# Genome-wide by environment interaction studies (GWEIS) of depressive symptoms and psychosocial stress in UK Biobank and Generation Scotland

**DOI:** 10.1101/479691

**Authors:** Aleix Arnau-Soler, Erin Macdonald-Dunlop, Mark J. Adams, Toni-Kim Clarke, Donald J. MacIntyre, Keith Milburn, Lauren Navrady, Generation Scotland, Major Depressive Disorder Working Group of the Psychiatric Genomics Consortium, Caroline Hayward, Andrew M. McIntosh, Pippa A. Thomson

## Abstract

Stress is associated with poorer physical and mental health. To improve our understanding of this link, we performed genome-wide association studies (GWAS) of depressive symptoms and genome-wide by environment interaction studies (GWEIS) of depressive symptoms and stressful life events (SLE) in two UK population cohorts (Generation Scotland and UK Biobank). No SNP was individually significant in either GWAS, but gene-based tests identified six genes associated with depressive symptoms in UK Biobank (*DCC, ACSS3, DRD2, STAG1, FOXP2 and KYNU; p* < 2.77×10^-6^). Two SNPs with genome-wide significant GxE effects were identified by GWEIS in Generation Scotland: rs12789145 (53kb downstream *PIWIL4; p =* 4.95×10^-9^; total SLE) and rs17070072 (intronic to *ZCCHC2; p =* 1.46×10^-8^; dependent SLE). A third locus upstream *CYLC2* (rs12000047 and rs12005200, *p* < 2.00×10^-8^; dependent SLE) when the joint effect of the SNP main and GxE effects was considered. GWEIS gene-based tests identified: *MTNR1B* with GxE effect with dependent SLE in Generation Scotland; and *PHF2* with the joint effect in UK Biobank (*p* < 2.77×10^-6^). Polygenic risk scores (PRS) analyses incorporating GxE effects improved the prediction of depressive symptom scores, when using weights derived from either the UK Biobank GWAS of depressive symptoms (*p =* 0.01) or the PGC GWAS of major depressive disorder (*p =* 5.91×10^-3^). Using an independent sample, PRS derived using GWEIS GxE effects provided evidence of shared aetiologies between depressive symptoms and schizotypal personality, heart disease and COPD. Further such studies are required and may result in improved treatments for depression and other stress-related conditions.

## INTRODUCTION

Mental illness results from the interplay between genetic susceptibility and environmental risk factors^1,2^. Previous studies have shown that the effects of environmental factors on traits may be partially heritable^3^ and moderated by genetics^4,5^. Major depressive disorder (MDD) is the most common psychiatric disorder with a lifetime prevalence of approximately 14% globally^6^ and with a heritability of approximately 37%^7^. There is strong evidence for the role of stressful life events (SLE) as risk factor and trigger for depression^8-12^. Genetic control of sensitivity to stress may vary between individuals, resulting in individual differences in the depressogenic effects of SLE, i.e., genotype-by-environment interaction (GxE)^4,13-16^. Significant evidence of GxE has been reported for common respiratory diseases and some forms of cancer^17-22^, and GxE studies have identified genetic risk variants not found by genome-wide association studies (GWAS)^23-27^.

Interaction between polygenic risk of MDD and recent SLE are reported to increase liability to depressive symptoms^4,16^; validating the implementation of genome-wide approaches to study GxE in depression. Most GxE studies for MDD have been conducted on candidate genes, or using polygenic approaches to a wide range of environmental risk factors, with some contradictory findings^28-32^. Incorporating knowledge about recent SLE into GWAS may improve our ability to detect risk variants in depression otherwise missed in GWAS^33^. To date, four studies have performed genome-wide by environment interaction studies (GWEIS) of MDD and SLE^34–37^, but this is the first study to perform GWEIS of depressive symptoms using adult SLE in cohorts of relatively homogeneous European ancestry.

Interpretation of GxE effects may be hindered by gene-environment correlation. Gene-environment correlation denotes a genetic mediation of associations through genetic influences on exposure to, or reporting of, environments^2,38^. Genetic factors predisposing to MDD may contribute to exposure and/or reporting of SLE^39^. To tackle this limitation, measures of SLE can be broken down into SLE likely to be independent of a respondent’s own behaviour and symptoms, or into dependent SLE, in which participants may played an active role exposure to SLE^40,41^. Different genetic influences with a higher heritability for reported dependent SLE than independent SLE^39,42-45^ suggest whereas GxE driven by independent SLE is likely to reflect a genetic moderation of associations between SLE and depression, GxE driven by dependent SLE may result from a genetic mediation of the association through genetically driven personality or behavioural traits. To test this we analysed dependent and independent SLE scores separately in Generation Scotland.

Stress contributes to many human conditions, with evidence of genetic vulnerability to the effect of SLE^46^. Therefore, genetic stress-response factors in MDD may also underlie the aetiology of other stress-linked disorders, with which MDD is often co-morbid^47,48^ (e.g. cardiovascular diseases^49^, diabetes,^50^ chronic pain^51^ and inflamation^52^). We tested the hypothesis that pleiotropy and shared aetiology between mental and physical health conditions may be due in part to genetic variants underlying SLE effects in depression.

In this study we conduct GWEIS of depressive symptoms incorporating data on SLE in two independent UK-based cohorts. We aimed to: i) identify loci associated with depressive symptoms through genetic response to SLE; ii) study dependent and independent SLE to support a contribution of genetically mediated exposure to stress; iii) assess whether GxE effects improve the proportion of phenotypic variance in depressive symptoms explained by genetic additive main effects alone; and iv) test for a significant overlap in the genetic aetiology of the response to SLE and mental and physical stress-related phenotypes.

## MATERIALS & METHODS

The core workflow of this study is summarized at Figure 1.

**Figure 1.**
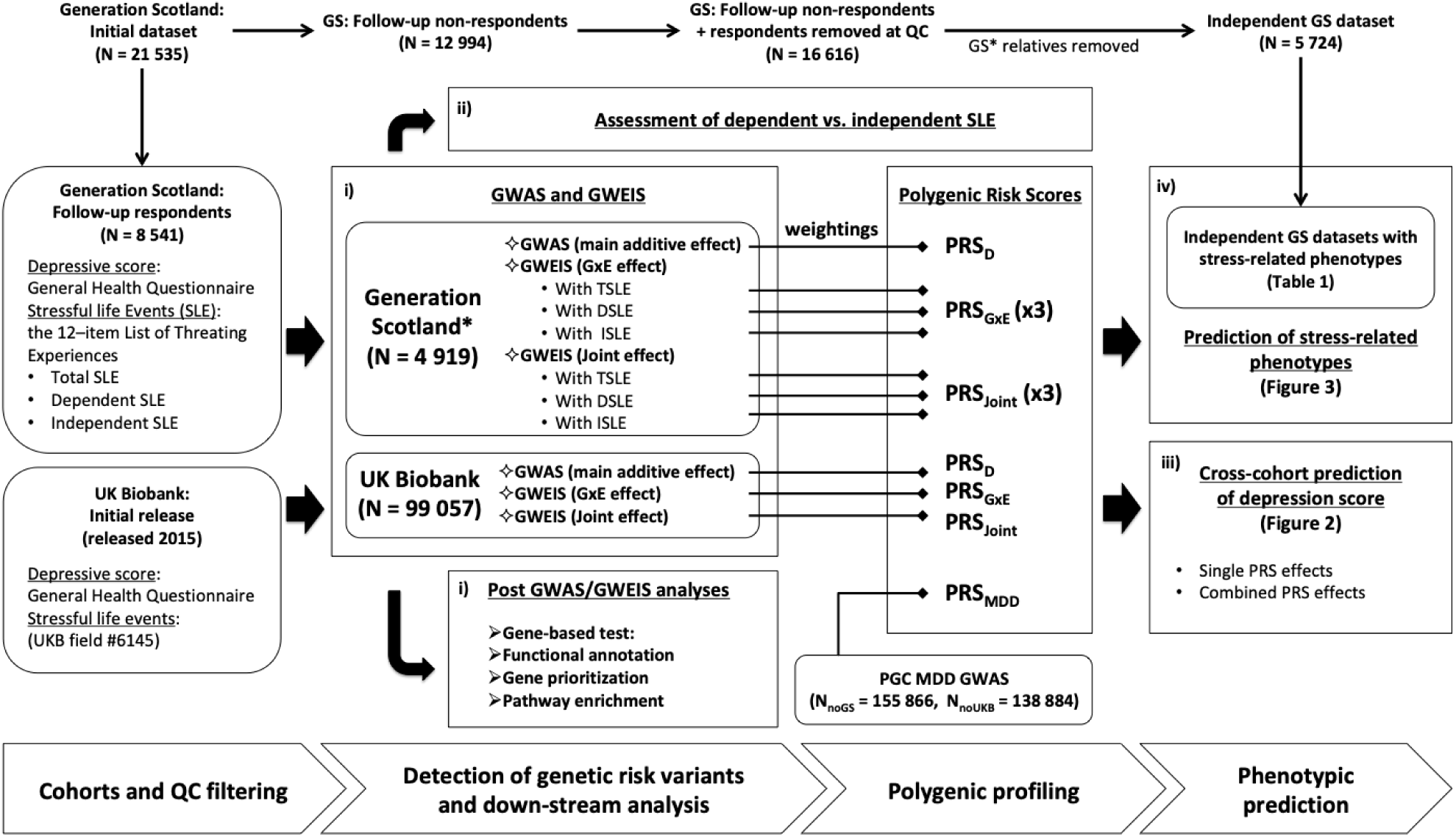
Study flowchart. Overview of analysis conducted on this study to achieve our aims: i) identify loci associated with depressive symptoms through genetic response to SLE; ii) test whether results of studying dependent and independent SLE support a contribution of genetically mediated exposure to stress; iii) assess whether GxE effects improve the proportion of phenotypic variance in depressive symptoms explained by genetic additive main effects alone and iv) test whether there is significant overlap in the genetic aetiology of the response to SLE and mental and physical stress-related phenotypes. Two core cohorts are used, Generation Scotland (GS) and UK Biobank (UKB). Summary statistics from Genome-Wide Association Studies (GWAS) and Genome-Wide by Environment Interaction Studies (GWEIS) are used to generate Polygenic Risk Scores (PRS). Summary statistics from Psychiatric Genetic Consortium (PGC) Major Depressive Disorder (MDD) GWAS are also used to generate PRS (PRS_MDD_). PRS weighted by: additive effects (PRS_D_ and PRS_MDD_), GxE effects (PRS_GXE_) and joint effects (the combined additive and GxE effect; PRS_Joint_), are used for phenotypic prediction. TSLE stands for Total number of SLE reported. DSLE stands for SLE dependent on an indivisual’s own behaviour. Coversely, ISLE stands for independent SLE. N stands for sample size. N_noGs_ stands for sample size with GS individuals removed. N_noUKB_ stands for sample size with UKB individuals removed.

## COHORT DESCRIPTIONS

### Generation Scotland (GS)

Generation Scotland is a family-based population cohort representative of the Scottish population^53^. At baseline, blood and salivary DNA samples were collected, stored and genotyped at the Wellcome Trust Clinical Research Facility, Edinburgh. Genome-wide genotype data was generated using the lllumina HumanOmniExpressExome-8 vl.O DNA Analysis BeadChip (San Diego, CA, USA) and Infinium chemistry^54^. The procedures and details for DNA extraction and genotyping have been extensively described elsewhere^55,56^. 21 525 participants were re-contacted to participate in a follow-up mental health study (Stratifying Resilience and Depression Longitudinally, STRADL), of which 8 541 participants responded providing updated measures in psychiatric symptoms and SLE through self-reported mental health questionnaires^57^. Samples were excluded if: they were duplicate samples, had diagnoses of bipolar disorder, no SLE data (non-respondents), were population outliers (mainly non-Caucasians and Italian ancestry subgroup), had sex mismatches, or were missing more than 2% of genotypes. SNPs were excluded if: missing more than 2% of genotypes, Hardy-Weinberg Equilibrium test *p <* lxlCf^6^, or minor allele frequency less than 1%. Further details of the GS and STRADL cohort are available elsewhere^53,57-59^. All components of GS and STRADL obtained ethical approval from the Tayside Committee on Medical Research Ethics on behalf of the NHS (reference 05/s1401/89). After quality control, individuals were filtered by degree of relatedness (pi-hat < 0.05), maximizing retention of those individuals reporting a higher number of SLE. The final dataset comprised data on 4 919 unrelated individuals (1 929 men; 2 990 women) and 560 351 SNPs.

### Independent GS datasets

Additional datasets for a range of stress-linked medical conditions and personality traits were created from GS (N = 21 525) excluding respondents and their relatives (N = 5 724). Following the same quality control criteria detailed above, we maximized unrelated non-respondents for retention of cases, or proxy cases (see below), to maximize the information available for each phenotype. This resulted in independent datasets with unrelated individuals for each trait. Differences between respondents and non-respondents are noted in the figure legend of Table 1.

**Table 1.** GS samples with stress-related phenotypes.

| **Trait** | **N** | **Males/Females** | **N SNPs** | **N Cases** | **N Controls** |
| --- | --- | --- | --- | --- | --- |
| Alzheimer (R) | 3377 | 1475/1903 | 560622 | **655** | **2722** |
| Asthma | 3390 | 1500/1890 | 560569 | **555** | **2835** |
| Asthma (R) | 3375 | 1470/1905 | 560432 | **910** | **2465** |
| Bowel cancer (R) | 3386 | 1495/1891 | 560630 | **672** | **2714** |
| Breast cancer | 3388 | 1486/1902 | 560611 | **83** | **3305** |
| Breast cancer (R) | 3386 | 1482/1904 | 560579 | **564** | **2822** |
| Chronic obstructive pulmonary disease | 3387 | 1496/1891 | 560591 | **73** | **3314** |
| Chronic obstructive pulmonary disease (R) | 3387 | 1474/1913 | 560620 | **553** | **2834** |
| Depression | 3385 | 1495/1890 | 560584 | **483** | **2902** |
| Depression (R) | 3382 | 1506/1876 | 560514 | **731** | **2651** |
| Diabetes | 3388 | 1497/1891 | 560469 | **185** | **3203** |
| Diabetes (R) | 3389 | 1481/1908 | 560584 | **1144** | **2245** |
| Heart disease | 3392 | 1504/1888 | 560526 | **212** | **3180** |
| Heart disease (R) | 3377 | 1483/1894 | 560479 | **2254** | **1123** |
| High blood pressure | 3402 | 1501/1901 | 560508 | **729** | **2673** |
| High blood pressure (R) | 3372 | 1464/1908 | 560569 | **1901** | **1471** |
| Hip fracture (R) | 3388 | 1489/1899 | 560572 | **421** | **2967** |
| Lung cancer (R) | 3379 | 1492/1887 | 560600 | **798** | **2581** |
| Osteoarthritis | 3395 | 1486/1909 | 560640 | **411** | 2984 |
| Osteoarthritis (R) | 3383 | 1466/1917 | 560516 | **961** | **2422** |
| Parkinson (R) | 3388 | 1488/1900 | 560590 | **236** | **3152** |
| Prostate cancer (R) | 3381 | 1495/1886 | 560570 | **329** | **3052** |
| Rheumatoid arthritis | 3387 | 1490/1897 | 560618 | **93** | **3294** |
| Rheumatoid arthritis (R) | 3380 | 1487/1893 | 560543 | **765** | **2615** |
| Stroke | 3387 | 1492/1895 | 560613 | **81** | **3306** |
| Stroke (R) | 3385 | 1463/1922 | 560478 | **1506** | **1879** |
| Neuroticism* | 3421 | 1521/1900 | 560484 | - | - |
| Extraversion* | 3420 | 1520/1900 | 560476 | - | - |
| Schizotypal personality* | 2386 | 1065/1321 | 560369 | - | - |
| Mood disorder* | 2307 | 1040/1267 | 560318 | - | - |

Samples were maximized for retention of cases to maximize the information available for each trait. There was no preferential selection of relatives in pairs for quantitative phenotypes, in order to retain the underlying distribution. All individuals involved in the datasets listed above were non-respondents to the GS follow-up study. Compared to individuals included at GS GWEIS (respondents in GS follow-up), non-respondents were significantly: younger, from more socioeconomically deprived areas, generally less healthier and wealthier. Non-respondents were more likely to smoke, and less likely to drink alcohol, although they consumed more units per week, compared with respondents. At GS baseline, non-respondents experienced more psychological distress and reported higher scores in symptoms of GHQ-depression and GHQ-anxiety than respondents<sup>57</sup>.
The total target sample size (N), number of males and females in N, number of SNPs (N SNPs) in target sample size N: the number of SNPs used as predictors after clumping step range between 90650 - 91000. The number of cases and controls in the independent target sample is indicated for binary phenotypes only. Samples were mapping by proxy approach was used (i.e. where first-degree relatives of individuals with the disease were considered proxy cases and included into the group of cases) are indicated by (R). \*Assessed through self-reported questionnaires.

### UK Biobank (UKB)

This study used data from 99 057 unrelated individuals (47 558 men; 51 499 women) from the initial release of UKB genotyped data (released 2015; under UK Biobank project 4844.). Briefly, participants were removed based on UKB genomic analysis exclusion, non-white British ancestry, high missingness, genetic relatedness (kinship coefficient > 0.0442), QC failure in UK BiLEVE study, and gender mismatch. GS participants and their relatives were excluded and GS SNPs imputed to a reference set combining the UK10K haplotype and 1000 Genomes Phase 3 reference panels^60^. After quality control, 1 009 208 SNPs remained. UK Biobank received ethical approval from the NHS National Research Ethics Service North West (reference: 11/NW/0382). Further details on UKB cohort description, genotyping, imputation and quality control are available elsewhere^61-63^.

All participants provided informed consent.

## PHENOTYPE ASSESSMENT

### Stressful life events (SLE)

GS participants reported SLE experienced over the preceding 6 months through a self-reported brief life events questionnaire based on the 12-item List of Threating Experiences^40,64,65^ (Supplementary Table 1a). The total number of SLE reported (TSLE) consisted of the number of ‘yes’ responses. TSLE were subdivided into SLE potentially dependent or secondary to an individual’s own behaviour (DSLE, questions 6-11 in Supplementary Table 1a), and independent SLE (ISLE, questions 1-5 in Supplementary Table 1a; pregnancy item removed) following Brugha *et al*.^40,41^. Thus, 3 SLE measures (TSLE, DSLE and ISLE) were constructed for GS. UKB participants were screened for *“illness, injury, bereavement and stress”* (Supplementary Table 1b) over the previous 2 years using 6 items included in the UKB Touchscreen questionnaire. A score reflecting SLE reported in UKB (TSLEukb) was constructed by summing the number of ‘yes’ responses.

### Psychological assessment

GS participants reported whether their current mental state over the preceding 2 weeks differed from their typical state using a self-administered 28-item scaled version of The General Health Questionnaire (GHQ)^66-68^. Participants rated the degree and severity of their current symptoms with a four-point Likert scale (following Goldberg et al., 1997^68^). A final log-transformed GHQ was used to detect altered psychopathology and thus, assess depressive symptoms as results of SLE. In UKB participants, current depressive symptoms over the preceding 2 weeks were evaluated using 4 psychometric screening items (Supplementary Table 2), including two validated and reliable questions for screening depression^69^, from the Patient Health Questionnaire (PHQ) validated to screen mental illness^70,71^. Each question was rated in a four-point Likert scale to assess impairment/severity of symptoms. Due to its skewed distribution, a four-point PHQ score was formed from PHQ (0 = 0; 1 = 1-2; 2 = 3-5; 3 = 6 or more) to create a more normal distribution.

### Stress-related traits

Targeted GS stress-related phenotypes and sample sizes are shown in Table 1 and detailed elsewhere^53^. These conditions were selected from literature review based on previous evidence of a link with stress^46^ (see also Supplementary Material: third section). Furthermore, we created additional independent samples using mapping by proxy, where individuals with a self-reported first-degree relative with a selected phenotype were included as proxy cases. This approach provides greater power to detect susceptibility variants in traits with low prevalence^72^.

## STATISTICAL ANALYSES

### SNP-heritability and genetic correlation

Restricted maximum likelihood approach was applied to estimate SNP-heritability (h^2^_SNP_) of depressive symptoms and self-reported SLE measures, and within samples bivariate genetic correlation between depressive symptoms and SLE measures using GCTA^73^.

### GWAS analyses

GWAS were conducted in PUNK^74^. In GS, age, sex and 20 principal components (PCs) were fitted as covariates. In UKB, age, sex, and 15 PCs recommended by UKB were fitted as covariates. The genome-wide significance threshold was *p =* 5×10^-8^.

### GWEIS analyses

GWEIS were conducted on GHQ (the dependent variable) for TSLE, DSLE and ISLE in GS and on PHQ for TSLE_UKB_ in UKB fitting the same covariates detailed above to reduce error variance. GWEIS were conducted using an R plugin for PUNK^74^ developed by Almli *et al*.^75^ (https://epstein-software.github.io/robust-joint-interaction). This method implements a robust test, that jointly considers SNP and SNP-environment interaction effects from a full model (*Y*∼ *β*_0_ + *βSNP* + *βSLE* + *βSNPxSLE* + *βCovariates*) against a null model where both the SNP and SNPxSLE effects equal 0, to assess the joint effect (the combined additive main and GxE genetic effect at a SNP) using a nonlinear statistical approach that applies Huber-White estimates of variance to correct possible inflation due to heteroscedasticity (unequal variances across exposure levels). This robust test should reduce confounding due to differences in variance induced by covariate interaction effects^76^ if present. Additional code was added (courtesy of Prof. Michael Epstein^75^; Supplementary Material) to generate beta-coefficients and the p-value of the GxE term alone. In UKB, correcting for 1 009 208 SNPs and 1 exposure we established a Bonferroni-adjusted threshold for significance at *p =* 2.47×10^-8^ for both joint and GxE effects. In GS, correcting for 560 351 SNPs and 3 measures of SLE we established a genome-wide significance threshold of *p =* 2.97×10^-8^.

### Post-GWAS/GWEIS analyses

GWAS and GWEIS summary statistics were analysed using FUMA^77^ including: gene-based tests, functional annotation, gene prioritization and pathway enrichment (Supplementary Material).

### Polygenic profiling & prediction

Polygenic risk scores (PRS) weighting by GxE effects (PRS_GxE_) were generated using PRSice-2^78^ (Supplementary Material) in GS using GxE effects from UKB-GWEIS. In UKB, PRS_GxE_ were constructed using GxE effects from all three GS-GWEIS (TSLE, DSLE and ISLE as exposures) independently. PRS were also weighted in both samples using either UKB-GWAS or GS-GWAS statistics (PRS_D_), and summary statistics from Psychiatric Genetic Consortium (PGC) MDD-GWAS (released 2016; PRS_MDD_) that excluded GS and UKB individuals when required (N_noGs_ = 155 866; N_noUKB_ = 138 884). Furthermore, we calculated PRS weighted by the joint effects (the combined additive main and GxE genetic effects; PRS_Joint_) from either the UKB-GWEIS or GS-GWEIS. PRS predictions of depressive symptoms were permuted 10 000 times. Multiple regression models fitting PRS_GXE_ and PRS_MDD_, and both PRS_GXE_ and PRS_D_ were tested. All models were adjusted by same covariates used in GWAS/GWEIS. Null models were estimated from the direct effects of covariates alone. The predictive improvement of combining PRS_GXE_ and PRS_MDD_/PRS_D_ effects over PRS_MDD_/PRS_D_ effect alone was tested for significance using the likelihood-ratio test (LRT).

Prediction of PRS_D_, PRS_GXE_ and PRS_Joint_ on stress-linked traits were adjusted by age, sex and 20 PCs; and permuted 10 000 times. Empirical-*p*-values after permutations were further adjusted by false discovery rate (conservative threshold at *Empirical-p =* 6.16×10^-3^). The predictive improvement of fitting PRS_GXE_ combined with PRS_D_ and covariates over prediction of a phenotype using the PRS_D_ effect alone with covariates was assessed using LRT, and *LRT-p-v*alues adjusted by FDR (conservative threshold at *LRT-p =* 8.35×10^-4^).

## RESULTS

### Phenotypic and genetic correlations

Depressive symptoms scores and SLE measures were positively correlated in both UKB (r^2^ = 0.22, *p* < 2.2×10^-16^) and GS (TSLE-r^2^ = 0.21, *p* =1.69×10^-52^; DSLE-r^2^ = 0.21, *p =* 8.59×10^-51^; ISLE-r^2^ = 0.17, *p =* 2.33×10^-33^). Significant bivariate genetic correlation between depression and SLE scores was identified in UKB (rG = 0.72; *p* < 1×10’^5^, N = 50 000), but not in GS (rG = 1, *p* ≥ 0.056, N = 4 919; Supplementary Table 3a).

### SNP-heritability

(h^2^_SNP_). In UKB, a significant h^2^_SNP_ of PHQ was identified (h^2^_SNP_ = 0.090; *p* < 0.001; N = 99 057). This estimate remained significant after adjusting by TSLE_ukB_ effect (h^2^_SNP_ = 0.079; *p* < 0.001), suggesting a genetic contribution unique of depressive symptoms. The h^2^_SNP_ of TSLEukb was also significant (h^2^_SNP_ = 0.040, *p <* 0.001; Supplementary Table 3b). In GS, h^2^_SNP_ was not significant for GHQ (h^2^_SNP_ = 0.071, *p =* 0.165; N = 4 919). However, in an ad hoc estimation from the baseline sample of 6 751 unrelated GS participants (details in Supplementary Table 3b) we detected a significant h^2^_SNP_ for GHQ (h^2^_SNP_ = 0.135; *p* < 5.15×10^-3^), suggesting that the power to estimate h^2^_SNP_ in GS may be limited by sample size. Estimates were not significant for neither TSLE (h^2^_SNP_ = 0.061, *p =* 0.189; Supplementary Table 3b) nor ISLE (h^2^_SNP_ = 0.000, *p =* 0.5), but h^2^_SNP_ was significant for DSLE (h^2^_SNP_ = 0.131, *p =* 0.029), supporting a potential genetic mediation and gene-environment correlation.

### GWAS of depressive symptoms

No genome-wide significant SNPs were detected by GWAS in either cohort. Top findings *(p <* 1×10^-5^) are summarized in Supplementary Table 4. Manhattan and QQ plots are shown in Supplementary Figures 1-4. There was no evidence of genomic inflation (all λ _1000_ < 1.01).

### Post-GWAS analyses

Gene-based test identified six genes associated with PHQ using UKB-GWAS statistics at genome-wide significance (Bonferroni-corrected *p =* 2.77×10^−6^; *DCC, p =* 7.53×10^-8^; *ACSS3, p =* 6.51×10^-7^; *DRD2, p =* 6.55×10^-7^; *STAG1, p =* 1.63×10-^6^; *FOXP2, p =* 2.09×10^-6^; *KYNU, p =* 2.24×10^-6^; Supplementary Figure 8). Prioritized genes based on position, eQTL and chromatin interaction mapping are detailed in Supplementary Table 5. No genes were detected in GS-GWAS gene-based test (Supplementary Figures 9). No tissue enrichment was detected from GWAS in either cohort. Significant gene-sets and GWAS catalog associations for UKB-GWAS are reported in Supplementary Table 6. These included the *biological process:* positive regulation of long term synaptic potentiation, and *GWAS catalog associations:* brain structure, schizophrenia, response to amphetamines, age-related cataracts (age at onset), fibrinogen, acne (severe), fibrinogen levels, and educational attainment; all adjusted-*p* < 0.01. There was no significant gene-set enrichment from GS-GWAS.

### GWEIS of depressive symptoms

Manhattan and QQ plots are shown in Supplementary Figures 1-4. There was no evidence of GWEIS inflation for either UKB or GS (all λ _1000_ < 1.01). No genome-wide significant GWEIS associations were detected for SLE in UKB. GS-GWEIS using TSLE identified a significant GxE effect (*p* < 2.97×10^-8^) at an intragenic SNP on chromosome 11 (rs12789145, *p =* 4.95×10^-9^, [3 = 0.06, closest gene: *PIWIL4;* Supplementary Figure 5), and using DSLE at an intronic SNP in *ZCCHC2* on chromosome 18 (rs17070072, *p =* 1.46×10^-8^, 3 = −0.08; Supplementary Figure 6). In their corresponding joint effect tests both rs12789145 (*p =* 2.77×10^-8^) and rs17070072 *p =* 1.96×10^-8^) were significant. GWEIS for joint effect using DSLE identified two further significant SNPs in on chromosome 9 (rs12000047, *p =* 2.00×10^-8^, 3 = −0.23; rs12005200, *p =* 2.09×10^-8^, β = −0.23, LD *r*^*2*^*>* 0.8, closest gene: *CYLC2;* Supplementary Figure 7). None of these associations replicated in UKB *(p >* 0.05), although the effect direction was consistent between cohorts for the SNP close to *PIWL1* and SNPs at *CYLC2.* No SNP achieved genome-wide significant association in GS-GWEIS using ISLE as exposure. Top GWEIS results *(p* < l×10^-5^) are summarized in Supplementary Tables 7-10.

### Post-GWEIS analyses: gene-based tests

All results are shown in Supplementary Figures 10-17. Two genes were associated with PHQ using the joint effect from UKB-GWEIS (*ACSS3 p =* 1.61×10^-6^; *PHF2, p =* 2.28×10^-6^; Supplementary Figure 11). *ACSS3* was previously identified using the additive main effects, whereas *PHF2* was only significantly associated using the joint effects. Gene-based tests identified *MTNR1B* as significantly associated with GHQ on GS-GWEIS using DSLE in both GxE (*p =* 1.53×10^-6^) and joint effects (*p =* 2.38×10^-6^; Supplementary Figures 14-15).

### Post-GWEIS analyses: tissue enrichment

We prioritized genes based on position, eQTL and chromatin interaction mapping in brain tissues and regions. In UKB, prioritized genes with GxE effect were enriched for up-regulated differentially expressed genes from adrenal gland (adjusted-*p =* 3.58×10^-2^). Using joint effects, prioritized genes were enriched on up-regulated differentially expressed genes from artery tibial (adjusted-*p =* 4.34×10^-2^). In GS, prioritized genes were enriched: in up-regulated differentially expressed genes from artery coronary (adjusted-*p =* 4.55×10^-2^) using GxE effects with DSLE; in down-regulated differentially expressed genes from artery aorta tissue (adjusted-*p =* 4.71×10^-2^) using GxE effects with ISLE; in up-regulated differentially expressed genes from artery coronary (adjusted-*p =* 5.97×10^-3^, adjusted-*p =* 9.57×10^-3^) and artery tibial (adjusted-*p =* 1.05×10^-2^, adjusted-*p =* 1.55×10” ^2^) tissues using joint effects with both TSLE and DSLE; and in down-regulated differentially expressed genes from lung tissue (adjusted-*p =* 3.98×10^-2^) and in up- and down-regulated differentially expressed genes from the spleen (adjusted-*p =* 4.71×10^−2^) using joint effects with ISLE. There was no enrichment using GxE effect with TSLE.

### Post-GWEIS analyses: gene-sets enrichment

Significant gene-sets and GWAS catalog hits from GWEIS are detailed in Supplementary Tables 11-14, including for UKB *Biocarta:* GPCR pathway; *Reactome:* opioid signalling, neurotransmitter receptor binding and downstream transmission in the postsynaptic cell, transmission across chemical synapses, gastrin CREB signalling pathway via PKC and MAPK; *GWAS catalog:* post bronchodilator FEV1/FVC ratio, migraine and body mass index. In GS, enrichment was seen using TSLE and DLSE for *GWAS catalog:* age-related macular degeneration, myopia, urate levels and Heschl’s gyrus morphology; and using ISLE for *biological process:* regulation of transporter activity. All adjusted-*p* < 0.01.

### Cross-cohort prediction

In GS, PRS_D_ weighted by UKB-GWAS of PHQ significantly explained 0.56% of GHQ variance (*Empirical-p* < 1.10^-4^), similar to PRS_MDD_ weighted by PGC MDD-GWAS (R^2^ = 0.78%, *Empirical-p* < 1.10 ^4^). PRS_GxE_ weighted by UKB-GWEIS GxE effects explained 0.15% of GHQ variance (*Empirical-p =* 0.03, Supplementary Table 15). PRS_GxE_ fitted jointly with PRSmdd significantly improved prediction of GHQ (R^2^ = 0.93%, model *p =* 6.12×10^-11^; predictive improvement of 19%, *LRT-p =* 5.91×10^-3^) compared to PRSmdd alone. Similar to PRS_GxE_ with PRS_D_ (R^2^ = 0.69%, model *p =* 2.72×10^-8^; predictive improvement of 23%, *LRT-p =* 0.01). PRS_Joint_ weighted by UKB-GWEIS also predicted GHQ (R^2^ = 0.58%, *Empirical-p* < 1.10^-4^), although the variance explained was significantly reduced compared to the model fitting PRS_GxE_ and PRS_D_ together (*LRT-p =* 4.69×10^-7^), suggesting that additive and GxE effects should be modelled independently for polygenic approaches (Figure 2a).

**Figure 2.**
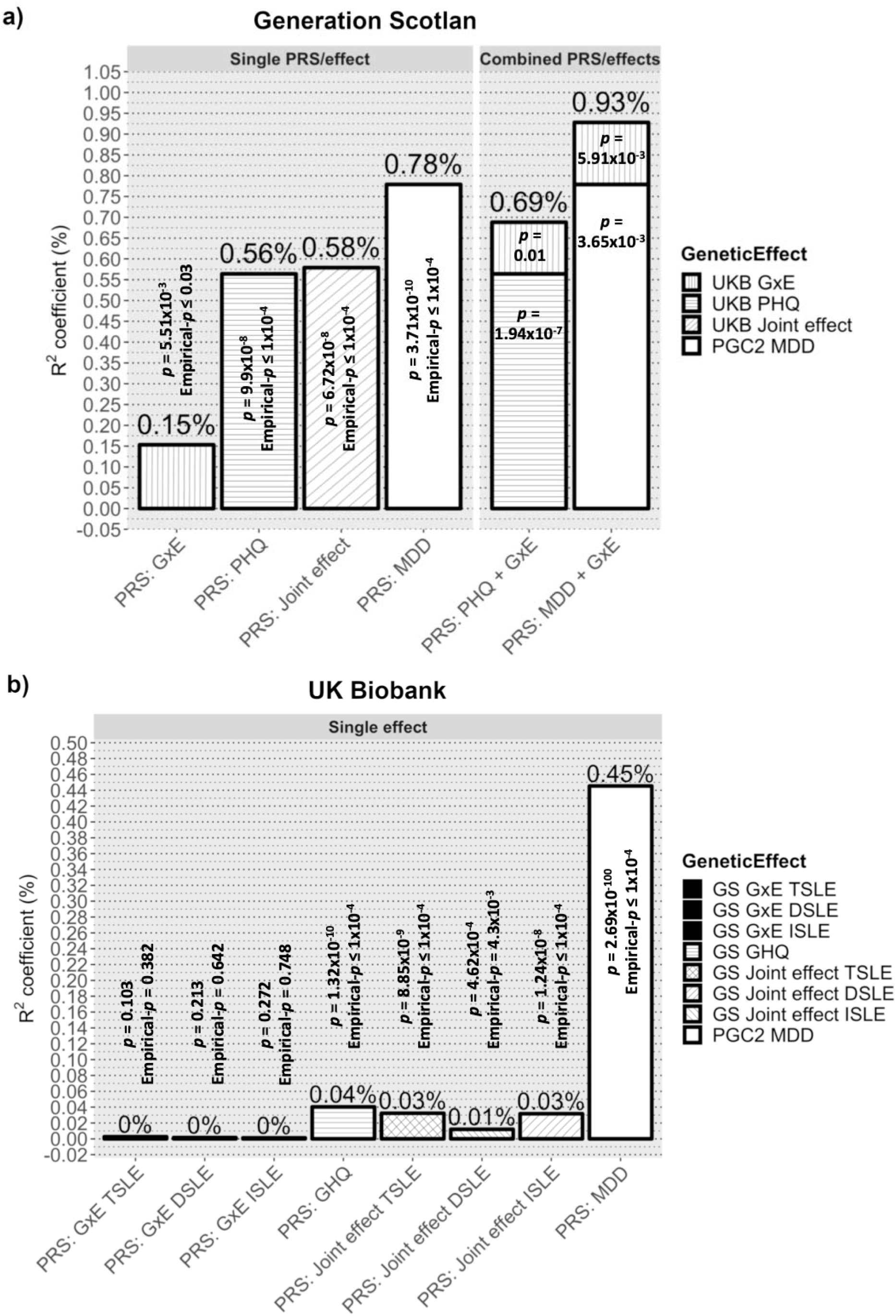
Prediction of depression scores by PRS_GXE_, PRS_D_, PRS_MDD_ & PRS_Joint_. Variance of depression score explained by PRS_GxE_ PRS_D_, PRS_MDD_ and PRS_Joint_ as single effect; and combining both PRSd and PRSmdd with PRS_GxE_ in single models. Prediction was conducted using **2a)** Generation Scotland and **2b)** UK Biobank as target sample. PRSgxe were weighted by cross sample GWEIS using GxE effect. PRS_D_ were weighted by cross sample GWAS of depressive symptoms effect. PRSmdd was weighted by PGC MDD-GWAS summary statistics. PRS_Joint_ were weighted by cross sample GWEIS using joint effect. A nominally significant gain in variance explained of GHQ of about 23% was seen in Generation Scotland when PRS_Gx_e was incorporated into a multiple regression model along with PRS_D_; and of about 19% when PRS_GxE_ was incorporated into a multiple regression model along with PRSmdd-Such gain was not seen in UK Biobank, but it must be noted that both PRS_D_ and PRS_MDD_ also explains much less variance of PHQ in UK Biobank than of GHQ in Generation Scotland. To note a noticeably reduction of variance explained by PRS_Joint_ compared to combined PRS/effects.

In UKB (Figure 2b), both PRS_D_ weighted by GS-GWAS of GHQ and PRS_MDD_ significantly explained 0.04% and 0.45% of PHQ variance, respectively (both *Empirical-p* < 1.10^-4^; Supplementary Table 15). PRS_GxE_ derived from GS-GWEIS GxE effect did not significantly predicted PHQ (TSLE-PRS_GxE_ *Empirical-p =* 0.382; DSLE-PRS_GxE_ *Empirical-p =* 0.642; ISLE-PRSqxe *Empirical-p =* 0.748). Predictive improvements by PRS_GxE_ effect fitted jointly with PRS_MDD_ or PRS_D_ were not significant (all *LRT-p* > 0.08). PRS_Joint_ significantly predicted PHQ (TSLE-PRS_Joint_: R^2^ = 0.032%, *Empirical-p* < 1.10^-4^; DSLE-PRSjoint: R^2^ = 0.012%, *Empirical-p =* 4.3×10^-3^; ISLE-PRS_Joint_: R^2^ = 0.032%, *Empirical-p <* 1.10^-4^), although the variances explained were significantly reduced compared to the models fitting PRS_GxE_ and PRS_D_ together (all *LRT-p* < 1.48×10^-3^).

### Prediction of stress-related traits

Prediction of stress-related traits in independent samples using PRS_D_, PRS_GxE_ and PRS_Joint_ are summarized in Figure 3a and Supplementary Table 16. Significant trait prediction after FDR adjustment (*Empirical-p* < 6.16×10^-3^, FDR-adjusted *Empirical-p* < 0.05) using both UKB and GS PRS_D_ was seen for: depression status, neuroticism and schizotypal personality. PRS_GxE_ weighted by GS-GWEIS GxE effect using ISLE significantly predicted depression status mapping by proxy (*Empirical-p =* 7.00×10^-4^, *FDR-adjusted Empirical-p =* 9.54×10^-3^).

**Figure 3.**
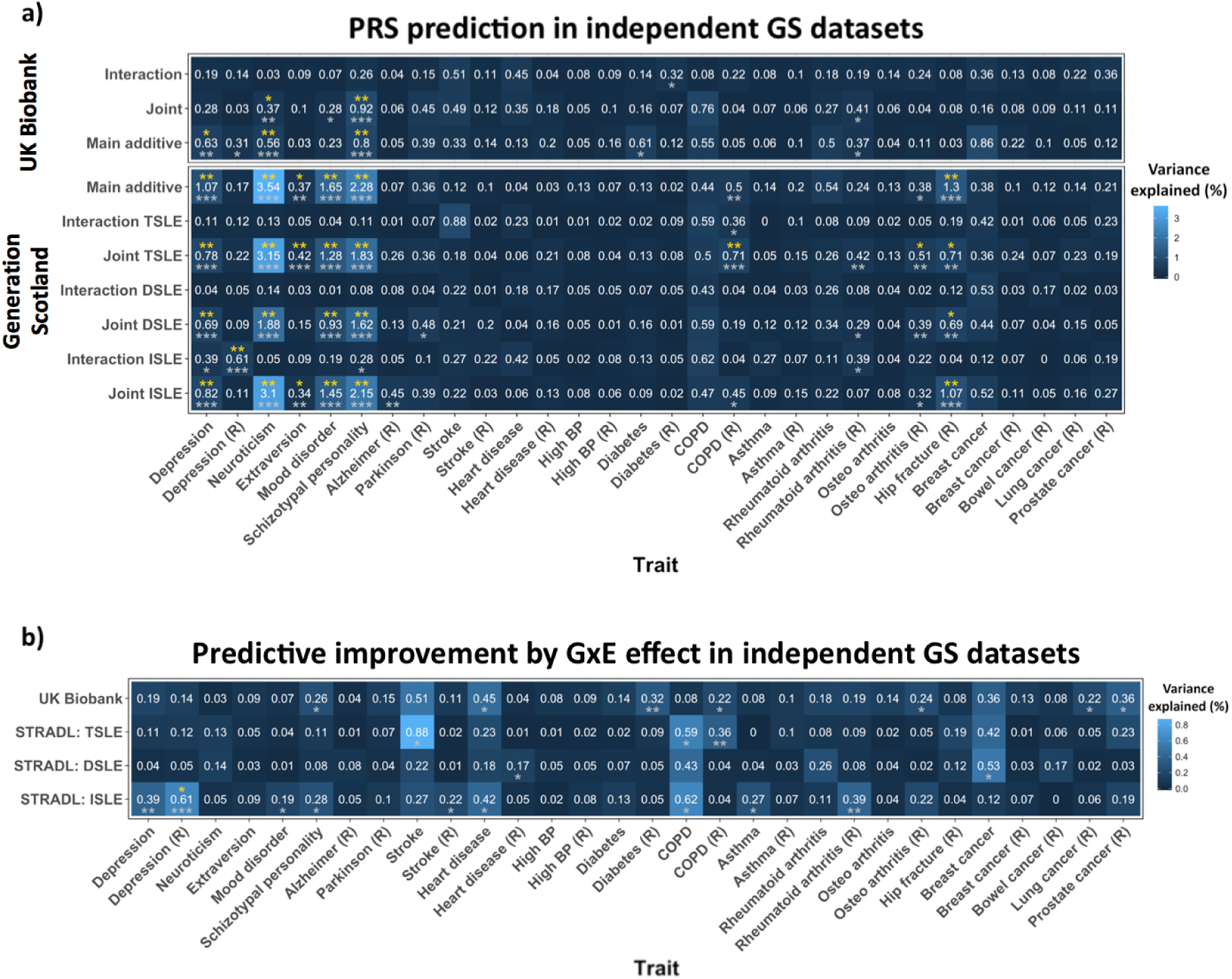
a) PRS prediction in independent GS datasets. Heatmap illustrating PRS prediction of a wide range of traits from GS listed in the x-axis (Table 1). (R) refers to traits using mapping by proxy approach (i.e. where first-degree relatives of individuals with the disease are considered proxy cases and included into the group of cases). Y-axis shows the discovery sample and the effect used to weight PRS. Numbers in cells indicate the % of variance explained, also represented by colour scale. Significance is represented by “*” according to the following significance codes: p-values **** < l×10 ^4^ < *** < 0.001 < ** < 0.01 * < 0.05; in grey *Empirical-p-values* after permutation (10 000 times) and in yellow *FDR-adjusted Empirical-p-values.* **b) Predictive improvement by GxE effect in independent GS datasets.** Heatmanp illustrating the predictive improvement as a result of incorporating PRS_GxE_ into a multiple model along with PRS_D_ and covariates (full model), over a model fitting PRS_D_ alone with covariates (null model); predicting a wide range of traits from GS listed in the x-axis (Table 1). Covariates: age, sex and 20 PCs. (R) refers to traits using mapping by proxy approach (i.e. where first-degree relatives of individuals with the disease are consider proxy cases and included into the group of cases). PRS_GxE_ are weighted by GWEIS using GxE effects. PRS_D_ were weighted by the GWAS of depressive symptoms additive main effects. The Y-axis shows the discovery sample used to weight PRS. Numbers in cells indicate the % of variance explained by the PRS_GxE_, also represented by colour scale. Notice that those correspond to the PRS_GxE_ predictions in Figure 2 when PRS_GxE_ are weighted by GxE effects. Significance was tested by Likelihood ratio tests (LRT): full model including PRS_D_ + PRS_GxE_ vs. null model with PRS_D_ alone (covariates adjusted). Significance is represented by “*” according to the following significance codes: p-values *** < 0.001 < ** < 0.01 * < 0.05; in grey *LRT-p-vales* and in yellow *FDR-adjusted LRT-p-values.*

Nominally significant predictive improvements (*LRT-p* < 0.05) of fitting PRS_GxE_ over the PRS_D_ effect alone using summary statistics generated from both UKB and GS were detected for schizotypal personality, heart diseases and COPD by proxy (Figure 3b). PRS_GxE_ weighted by GS-GWEIS GxE effect using ISLE significantly improved prediction over PRS_D_ effect alone of depression status mapping by proxy after FDR adjustment (*LRT-p =* 1.96×10^-4^, *FDR-adjusted LRT-p =* 2.35×10^-2^).

## DISCUSSION

This study performs GWAS and incorporates data on recent adult stressful life events (SLE) into GWEIS of depressive symptoms, identifies new loci and candidate genes for the modulation of genetic response to SLE; and provides insights to help disentangle the underlying aetiological mechanisms increasing genetic liability through SLE to both depressive symptoms and stress-related traits.

SNP-heritability of depressive symptoms (h^2^_SNP_ *=* 9-13%), were slightly higher than estimates from African American populations^34^, and over a third larger than estimates in MDD from European samples^79^. h^2^_SNP_ for PHQ in UKB (9.0%) remained significant after adjusting for SLE (7.9%). Thus, although some genetic contributions may be partially shared between depressive symptoms and reporting of SLE, there is still a relatively large genetic contribution unique to depressive symptoms. Significant h^2^_SNP_ of DSLE in GS (13%) and TSLE_UKB_ in UKB (4%), which is mainly composed of dependent SLE items, were detected similar to previous studies (8% and 29%)^34,43^. Conversely, there was no evidence for heritability of independent SLE. A significant bivariate genetic correlation between depressive symptoms and SLE (rG = 0.72) was detected in UKB after adjusting for covariates, suggesting that there are shared common variants underlying self-reported depressive symptoms and SLE. This bivariate genetic correlation was smaller than that estimated from African American populations (rG = 0.97; *p=* 0.04; N = 7 179)^34^. Genetic correlations between SLE measures and GHQ were not significant in GS (N = 4 919; rG = 1; all *p >* 0.056), perhaps due to a lack of power in this smaller sample.

Post-GWAS gene-based tests detected six genes significantly associated with PHQ (*DCC, ACSS3, DRD2, STAG1, FOXP2* and *KYNU).* Previous studies have implicated these genes in liability to depression (see Supplementary Table 17), and three of them are genome-wide significant in gene-based tests from the latest meta-analysis of major depression that incudes UKB (*DCC, p =* 2.57×10 ^-14^; *DRD2, p =* 5.35×10^-14^; and *KYNU, p =* 2.38×10^-6^; N = 807 553)^80^. This supports the implementation of quantitative measures such as PHQ to detect genes underlying lifetime depression status^81^. For example, significant gene ontology analysis of the UKB-GWAS identified enrichment for positive regulation of long-term synaptic potentiation, and for previous GWAS findings of brain structure^82^, schizophrenia^83^ and response to amphetamines^84^.

The key element of this study was to conduct GWEIS of depressive symptoms and recent SLE. We identified two loci with significant GxE effect in GS. However, none of these associations replicated in UKB (*p >* 0.05). The strongest association was using TSLE at 53kb down-stream of *PIWIL4* (rs12789145). PIWIL4 is brain-expressed and involved in chromatin-modification^85^, suggesting it may moderate the effects of stress on depression. It encodes HIWI2, a protein thought to regulate OTX2, which is critical for the development of forebrain and for coordinating critical periods of plasticity disrupting the integration of cortical circuits^86,87^. Indeed, an intronic SNP in *PIWIL4* was identified as the strongest GxE signal in ADHD using mother’s warmth as environmental exposure^88^. The other significant GxE identified in our study was in *ZCCHC2* using DSLE. This zinc finger protein is expressed in blood CD4+ T-cells and is down regulated in individuals with MDD^89^ and in those resistant to treatment with citalopram^90^. No GxE effect was seen using ISLE as exposure.

No significant locus or gene with GxE effect was detected in UKB-GWEIS. Nevertheless, joint effects (combined additive main and GxE genetic effects) identified two genes significantly associated with PHQ (*ACSS3* and *PHF2;* see Supplementary Table 17). *PHF2* was recently detected as genome-wide significant at the latest meta-analysis of depression^80^. Notably, *PHF2* paralogs have already been link with MDD through stress-response in three other studies^91-93^. Joint effects in GS also detected an additional significant association upstream *CYLC2*, a gene nominally associated (p < 1×10’^5^) with obsessive-compulsive disorder and Tourette’s Syndrome^94^. Gene-based test from GS-GWEIS identified a significant association with *MTNR1B*, a melatonin receptor gene, using DSLE (both GxE and joint effect; Supplementary Table 17). Prioritized genes using GxE effects were enriched in differentially expressed genes from several tissues including the adrenal gland, which releases cortisol into the blood-stream in response to stress, thus playing a key role in the stress-response system, reinforcing a potential role of GxE in stress-related conditions.

The different instruments and sample sizes available make it hard to compare results between cohorts. Whereas GS contains deeper phenotyping measurements of stress and depressive symptoms than UKB, the sample size is much smaller, which may be reflected in the statistical power required to detect reliable GxE effects. Furthermore, the presence and size of GxE are dependent on their parameterization (i.e. the measurement, scale and distribution of the instruments used to test such interaction)^95^. Thus, GxE may be incomparable across GWEIS due to differences in both phenotype assessment and stressors tested. Although our results suggest that both depressive symptom measures are correlated with lifetime depression status, different influences on depressive symptoms from the SLE covered across studies may contribute to lack of stronger replication. Instruments in GS cover a wider range of SLE and are more likely to capture changes in depressive symptoms as consequence of their short-term effects. Conversely, UKB could capture more marked long-term effects, as SLE were captured over 2 years compared to 6 months in GS. New mental health questionnaires covering a wide range of psychiatric symptoms and SLE in the last release of UKB data provides the opportunity to create more similar measures to GS in the near future. Further replication in independent studies with equivalent instruments is required to validate our GWEIS findings. Despite these limitations and a lack of overlap in the individual genes prioritised from the two GWEIS, replication was seen in the predictive improvement of using PRSGXE derived from the GWEIS GxE effects to predict stress-related phenotypes.

The third aim of this study was to test whether GxE effect could improve predictive genetic models, and thus help to explain deviation from additive models and missing heritability of MDD^96^. Multiple regression models suggested that inclusion of PRS_GXE_ weighted by GxE effects could improve prediction of an individual’s depressive symptoms over use of PRS_MDD_ or PRS_D_ weighted by additive effects alone. In GS, we detected a predictive gain of 19% over PRS_MDD_ weighted by PGC MDD-GWAS, and a gain of 23% over PRS_D_ weighted by UKB-GWAS (Figure 2a). However, these findings did not surpass stringent Bonferroni-correction and could not be validated in UKB. This may reflect in the poor predictive power of the PRS generated from the much smaller GS discovery sample. The results show a noticeably reduced prediction using PRS_Joint_ weighted by joint effects, which suggests that the genetic architecture of stress-response is at least partially independent and differs from genetic additive main effects. Therefore, our results from multiple regression models suggest that for polygenic approaches main and GxE effects should be modelled independently. SLE effects are not limited to mental illness^46^.

Our final aim was to investigate shared aetiology between GxE for depressive symptoms and stress-related traits. Despite the differences between the respondents and non-respondents (Table 1 legend), a significant improvement was seen predicting depressive status mapping by proxy cases using GxE effect from GS-GWEIS with independent SLE (*FDR-adjusted LRT-p* = 0.013), but not with dependent SLE. GxE effects using statistics generated from both discovery samples, despite the differences in measures, nominally improved the phenotypic prediction of schizotypal personality, heart disease and the proxy of COPD (*LRT-p* < 0.05). Other studies have found evidence supporting a link between stress and depression in these phenotypes that support our results (see Supplementary Material for extended review) and suggest, for instance, potential pleiotropy between schizotypal personality and stress-response. Our findings point to a potential genetic component underlying a stress-response-depression-comorbidities link due, at least in part, to shared stress-response mechanisms. A relationship between SLE, depression and coping strategies such as smoking suggests that perhaps, genetic stress-response may modulate adaptive behaviours such as smoking, fatty diet intake, alcohol consumption and substance abuse. This is discussed further in the Supplementary Material.

In this study, evidence for SNPs with significant GxE effects came primarily from the analyses of dependent SLE and not from independent SLE. This supports a genetic effect on probability of exposure to, or reporting of SLE, endorsing a gene-environment correlation. Chronic stress may influence cognition, decision-making and behaviour eventually leading to higher risk-taking^97^. These conditions may also increase sensitivity to stress amongst vulnerable individuals, including those with depression, who also have a higher propensity to report SLE, particularly dependent SLE^39^. A potential reporting bias in dependent SLE may be mediated as well by heritable behavioural, anxiety or psychological traits such as risk-taking^43,98^. Furthermore, individuals vulnerable to MDD may expose themselves into environments of higher risk and stress^14^. This complex interplay, reflected in the form of a gene-environment correlation effect, would hinder the interpretation of GxE effects from GWEIS as pure interactions. A mediation of associations between SLE and depressive symptoms through genetically driven sensitivity to stress, personality or behavioural traits would support the possibility of subtle genotype-by-genotype (GxG) interactions, or genotype-by-genotype-by-environment (GxGxE) interactions contributing to depression ‘. In contrast, PRS prediction of the stress-related traits: schizotypal personality, heart disease and COPD, was primarily from derived weights using independent SLE, suggesting that a common set of variants moderate the effects of SLE across stress-related traits and that larger sample sizes will be required to detect the individual SNPs contributing to this. Thus, our finding supports the inclusion of environmental information into GWAS to enhance the detection of relevant genes. Results of studying dependent and independent SLE support a contribution of genetically mediated exposure to and/or reporting of SLE, perhaps through sensitivity to stress as mediator.

This study emphasises the relevance of GxE in depression and human health in general and provides the basis for future lines of research.

## Supporting information

## ACKNOWLEDGMENTS

See Supplementary Material: acknowledgments.

## FINANCIAL DISCLOSURE

The authors declare no conflict of interest.

Supplementary Material is available at Translational Psychiatry’s website.

