## Supplementary material for "Genome-wide by environment interaction studies (GWEIS) of depressive symptoms and psychosocial stress in UK Biobank and Generation Scotland"

This document contains supplementary information for: ***Arnau-Soler et al*.** **Genome-wide by environment interaction studies (GWEIS) of depressive symptoms and psychosocial stress in UK Biobank and Generation Scotland.**

R code to perform GWEIS.	3

Statistical analyses – appendix.	4

Evidence supporting a link between stress and depression in stress-related phenotypes.	5

Major Depressive Disorder Working Group of the Psychiatric Genomics Consortium Authorship	14

Acknowledgments	21

Supplementary Figures	22

Supplementary Figure 1. Manhattan and QQ plots of Generation Scotland GWAS and GWEIS using TSLE as exposure	22

Supplementary Figure 2. Manhattan and QQ plots of Generation Scotland GWAS and GWEIS using DSLE as exposure	23

Supplementary Figure 3. Manhattan and QQ plots of Generation Scotland GWAS and GWEIS using ISLE as exposure	24

Supplementary Figure 4. Manhattan and QQ plots of UK Biobank GWAS and GWEIS	25

Supplementary Figure 5. GxE effect derived from GWEIS on Generation Scotland using TSLE as exposure, regional plot of locus around rs12789145 on chromosome 11	26

Supplementary Figure 6. GxE effect derived from GWEIS on Generation Scotland using DSLE as exposure, regional plot of locus around rs17070072 on chromosome 18	27

Supplementary Figure 7. Joint effect derived from GWEIS on Generation Scotland using DSLE as exposure, regional plot of locus around rs17070072 on chromosome 9	28

Supplementary Figure 8. Manhattan plot of gene-based test of PHQ UKB GWAS	29

Supplementary Figure 9. Manhattan plot of gene-based test of GHQ GS GWAS	30

Supplementary Figure 10. Manhattan plot of gene-based test of PHQ UKB GWEIS using TSLE_UKB_ as exposure for GxE effect	31

Supplementary Figure 11. Manhattan plot of gene-based test of PHQ UKB GWEIS using TSLE_UKB_ as exposure for joint effect	32

Supplementary Figure 12. Manhattan plot of gene-based test of GHQ GS GWEIS using TSLE as exposure for GxE effect	33

Supplementary Figure 13. Manhattan plot of gene-based test of GHQ GS GWEIS using TSLE as exposure for joint effect	34

Supplementary Figure 14. Manhattan plot of gene-based test of GHQ GS GWEIS using DSLE as exposure for GxE effect	35

Supplementary Figure 15. Manhattan plot of gene-based test of GHQ GS GWEIS using DSLE as exposure for joint effect	36

Supplementary Figure 16. Manhattan plot of gene-based test of GHQ GS GWEIS using ISLE as exposure for GxE effect	37

Supplementary Figure 17. Manhattan plot of gene-based test of GHQ GS GWEIS using ISLE as exposure for joint effect	38

### R code to perform GWEIS.

The R script we used to perform GWEIS is a modified version from the original R script developed by Almli *et al.*^1^ (<https://epstein-software.github.io/robust-joint-interaction>). The modifications added into the code output beta coefficients from both SNP and SNPxE terms and the p-value for just the interaction (GxE) term into the final GWEIS summary statistics.

Using the original R script (version 20 Apr 2017), a version of the script as the one we used can be achieved by replacing from the source code:

**r <- c(p_model,p_robust)**

with:

**wald_int_only<-((gee.fit$beta[3])^2)/gee.fit$vbeta[3,3]   # this is the robust wald test for just the interaction parameter only**

**p_int_only<-pchisq(wald_int_only,1,lower.tail=F)    # this is the p-value for the wald test**

**r <- c(p_model,p_robust,beta_est[c(2,3)],p_int_only) # The ‘beta_est’ vector contains all the regression coefficients for all parameters in the GEE model.**

These additional lines of code were courtesy of Prof. Michael Epstein^1^.

### Statistical analyses – appendix.

**Post-GWAS/GWEIS analyses.** GWAS and GWEIS (for both GxE and joint effects) summary statistics were analysed using FUMA^1^ (<http://fuma.ctglab.nl>). Gene-based tests were conducted using MAGMA^2^ through FUMA platform using the default MAGMA settings. Genome-wide significance at gene-based test was set at the Bonferroni-corrected significance threshold *p* = 0.05/18 068 = 2.77x10^-6^. FUMA was also used to assess functional annotation, gene prioritization and pathway enrichment using lead SNPs from associated genomic loci. SNPs were clumped according to linkage disequilibrium (r^2^ = 0.6) using default settings to identify independent lead SNPs with a *p* < 1x10^-5^. Gene mapping was performed in protein-coding genes under default settings using: positional mapping, eQTL mapping in 10 brain tissues from GTEx, and chromatin interaction mapping using Hi-C data from two brain regions: dorsolateral/prefrontal cortex and hippocampus. The MHC region (25-34 Mb) was excluded from these analyses. We assessed gene-set enrichment of: differentially expressed genes in 53 tissue types, Canonical Pathways, and Gene Ontology terms. Previously reported associations from the GWAS catalog^3^ were reported for independent lead SNPs and/or their proxy SNPs.

**Polygenic profiling & prediction.** Polygenic risk scores (PRS) were constructed using PRSice-2^4^ (clump-based pruning r^2^ = 0.1, 10MBp window) for thirteen *p* thresholds (<0.001, <0.005, <0.01, <0.02, <0.03, <0.04, <0.05, <0.1, <0.2, <0.3, <0.4, <0.5, <=1) and standardized to a mean of 0. We calculated the proportion of variance explained using R^2^ and Nagelkerke’s R^2^ coefficients for binary traits. Polygenic risk scores weighting by GxE effects (PRS_GxE_) were generated as standard PRS^5^ using summary statistics from GWEIS as input (discovery sample), instead of summary statistics from GWAS. Thus, weighting PRS by GxE effects (or joint effects) instead of main additive effects.

### Evidence supporting a link between stress and depression in stress-related phenotypes.

We hypothesized that pleiotropy between stress-related conditions, both mental and physical, may be due in part to shared genetic stress-response mechanisms. In this study we report evidence, albeit weak, for an effect of GxE involved in depressive symptoms into other stress-related traits, in particular for schizotypal personality, heart disease and chronic obstructive pulmonary disease (COPD). We believe that at least partially, genetics underlying GxE detected in GWEIS of depressive symptoms belongs to unique stress-response mechanisms also involved in the aetiology of these other phenotypes. There are many studies on the literature that provide evidence supporting a link between stress and depression in these phenotypes that supports our conclusion.

***Schizotypal personality.*** One of the main characteristics of schizotypal personality is an abnormal perception of experiences and dysfunctional coping strategies^6-8^, which overlaps with autistic personality traits, anxiety and depression^9-11^. Schizotypal personality correlates with SLE and negative coping strategies in healthy individuals^12^. The effect of stress on schizotypal personality traits is moderated by genetics^13^ and mediate by dysfunctional coping^12,13^. Individuals with high schizotypal personality have been shown to report increased subjective stress and to display blunted and delayed cortisol response after acute psychosocial stress, compared to individuals with low schizotypal personality, and have higher baseline cortisol levels, suggesting a limited physiologically adaptation to SLE and a feasible link with cardiovascular comorbidities through chronic overactivation of the sympathetic nervous system^14^.

***Heart disease.*** A direct relationship between the effects of psychological stress and cardiovascular disease has been extensively reported^15-21^. Depression, as well as anger and anxiety, may be a manifestation form underlying negative emotions, and it has been proposed that such underlying negativity may origin the link between these conditions and cardiovascular disease^22^. Psychosocial stress associated with anxiety disorders increase autonomic arousal via the hypothalamic-pituitary axis and circulating cathecolamines, which is associated to an elevated risk of pro-inflammatory state and hypertension leading to the development of coronary heart disease^23^. Previous studies also link anxiety disorder to cardiovascular disease^16^, and evidence suggests a potential role for inflammation as an underlying mechanism linking chronic stress and an associated increased risk of coronary heart disease^16,17,24^. Psychological stress is a risk factor of hypertension^18,25^, acute myocardial infarction^21^, pathogenesis of coronary artery disease^16^, cardiovascular morbidity and mortality^19^, within other cardiovascular disease, with a potential elevated risk in women^20^. A consistent relationship between the occurrence of major depression episodes and cardiovascular disease has been demonstrated, and evidence suggest that there is a continuum spectrum of risk for coronary artery disease associated with depression according to the magnitude of depressive symptoms^16^. Smoking is a coping strategy^26^ and behavioural risk factor associated with depression^27^, and risk of suffering a cardiac event is substantially higher in depressed individuals who smoke than in depressed individuals who do not smoke^28^.

***Chronic obstructive pulmonary disease (COPD).*** In this study, we suggested that GxE effect in depressive symptoms is related with risk of COPD. The prevalence of depressive symptoms, depression and risk of anxiety is higher among COPD patients^29^,^30^,^31^. SLE have been associated with increased depressive symptoms, anxiety and worse quality of life in individuals with COPD compared to healthy controls^32,33^, with significant SLE-by-COPD interaction suggesting a substantially greater detrimental effect of stress in those individuals suffering COPD^32^. Participants with COPD have reported to experience the same number and severity of non-illness-related occurrence of SLE and perceived stress as non-COPD healthy participants, suggesting that individuals with COPD may be more vulnerable to detrimental effects of SLE than healthy controls^32^. The experience of anxiety and distress produced by breathing difficulties is an intrinsic source of stress directly related to COPD^34,35^. However, poor coping skills may be the principal psychological problem that contributes to psychological distress and poor quality of life among COPD patients^36^. Depression and heart failure are associated with disease-specific health-related quality of life in COPD patients^37^. Smoking is recognized as the most important risk factor for COPD^38^. Lifetime cumulative stress, including psychological stress early in life, has been shown to have and additive effect over time on risk for heavy smoking^39^. Elevated risk for developing nicotine dependence has been shown in individuals who experienced trauma, and exacerbated by post-traumatic stress disorder^40^. Cigarette smoke disrupts homeostasis of the alveolar capillary unit and generates endogenous mediators of inflammation that could result in COPD^41^. The exposure to inhaled toxics from smoke activates a cellular integrated stress response implicated in pulmonary pathology and lung disease^42^. However, well-known behavioural risk factors such as smoking become extremely difficult to quit in those individuals with high life stress and depression levels^27,43^. In addition, although smokers often report smoking as a strategy to relieve stress, nicotine dependency is reported to exacerbate psychological stress, showing stress levels of smokers being higher than those of non-smokers and reporting worse mood, specially between cigarettes^44^.

Interestingly, a post hoc test conducted in our study showed that PRS weighted by GxE effect derived from Generation Scotland GWEIS using TSLE that predicts COPD also predicts number of cigarettes smoked in an average week in 75 smokers from an independent GS sample (*p* = 4.43x10^-6^). This was not replicated using the UKB GxE effect,. However, it suggests that genetic stress-response and stress-related traits may be partially related through GxE driven coping styles. Increased risk for smoking is also linked to other psychiatric disorders where smoking may play an important role either as a causal factor, as an agent promoting brain changes, or as modulator of medication’s effect^45^. Asthma, a separate respiratory disease has also shown robust relationships with psychological stress, depression, neuro-psychiatric factors and neuroendocrine, immunologic and inflammatory systems^24,46-50^.

***Other stress-related traits*.** Chronic psychological stress has been recognised to influence the immune system and to have an impact on many inflammatory process and disease, and dysregulated inflammatory processes haves been linked to major depressive disorder and its comorbidities (e.g. asthma, diabetes, cardiovascular disorders, stroke)^24,51^. Therefore, the list of stress-related conditions link to depression through genetic stress-response mechanisms may be larger, including other personality traits such as extraversion or some cancers, among others. Findings suggest that extraversion was improved by incorporating SLE GxE effects into main additive effect as a combined joint effect; prediction otherwise missed by any GxE effect or UKB main (depressive symptoms) additive effects alone. Extraversion mediates (both positive and negative) life events and resilience, low extraversion being associated with higher numbers of SLE^24^. High levels of extraversion, which involves energy, positive emotionality and dominance, is linked with stress and adversity coping strategies through positive affective style, and related to resilience and higher level of social interaction allowing an environment of positive networks of social support^52^. Indeed, individuals who are genetically predisposed to high level of extraversion my perceive (and report) life events as more controllable and positive; which in turn may lead to higher level of extraversion^53^. Low extraversion is associated with depression and depressive symptoms^54,55^. Another interesting case is the link between stress and cancer. Stress has been associated with cancer^56-59^. A recent study with lifetime exposure has reported that perceived stress, principally from workplace psychological stress over cumulative duration of more than 15 years, was significantly positively associated with greater odds of cancer (including lung, colon, rectum, stomach cancers and Non-Hodgkin lymphoma) among men^57^. Interesting, piRNAs and PIWI-like proteins, including HIWI2 encoded by *PIWIL4,* a gene involved in chromatin-modification^60^ and located at the locus with the strongest genome-wide association between GxE effect and depressive symptoms reported in this study has been suggested as biomarkers and potential therapeutic targets for breast cancer^61,62^ and gastric cancer^63^. The effects of stress may have a pronounced impact on epigenetic changes like chromatin or histone modifications. Epigenetic processes have been widely implicated in cancer^64,65^.

***In summary.*** As reviewed, psychological stress is recognized to play a role in the risk and prognosis of many chronic illnesses, as well as depressive symptoms, including evidence for other stress-related comorbidities of depression not detailed above such as diabetes^66-71^ and stroke^72-74^, among others^75^. Furthermore, stress may influence the extent to which individuals engage with behavioural risk factors as coping strategies; including smoking, fatty diet intake, excessive alcohol intake or substance abuse. GxE effects and genetic stress-perception, stress-exposure or stress-response may drive selection of these coping strategies and modulate coping skills and how we work through SLE, and negative emotions.

### Major Depressive Disorder Working Group of the Psychiatric Genomics Consortium Authorship

Naomi R Wray* 1, 2

Stephan Ripke* 3, 4, 5

Manuel Mattheisen* 6, 7, 8, 9

Maciej Trzaskowski* 1

Enda M Byrne 1

Abdel Abdellaoui 10

Mark J Adams 11

Esben Agerbo 9, 12, 13

Tracy M Air 14

Till F M Andlauer 15, 16

Silviu-Alin Bacanu 17

Marie Bækvad-Hansen 9, 18

Aartjan T F Beekman 19

Tim B Bigdeli 17, 20

Elisabeth B Binder 15, 21

Douglas H R Blackwood 11

Julien Bryois 22

Henriette N Buttenschøn 8, 9, 23

Jonas Bybjerg-Grauholm 9, 18

Na Cai 24, 25

Enrique Castelao 26

Jane Hvarregaard Christensen 7, 8, 9

Toni-Kim Clarke 11

Jonathan R I Coleman 27

Lucía Colodro-Conde 28

Baptiste Couvy-Duchesne 2, 29

Nick Craddock 30

Gregory E Crawford 31, 32

Gail Davies 33

Ian J Deary 33

Franziska Degenhardt 34, 35

Eske M Derks 28

Nese Direk 36, 37

Conor V Dolan 10

Erin C Dunn 38, 39, 40

Thalia C Eley 27

Valentina Escott-Price 41

Farnush Farhadi Hassan Kiadeh 42

Hilary K Finucane 43, 44

Jerome C Foo 45

Andreas J Forstner 34, 35, 46, 47

Josef Frank 45

Héléna A Gaspar 27

Michael Gill 48

Fernando S Goes 49

Scott D Gordon 28

Jakob Grove 7, 8, 9, 50

Lynsey S Hall 11, 51

Christine Søholm Hansen 9, 18

Thomas F Hansen 52, 53, 54

Stefan Herms 34, 35, 47

Ian B Hickie 55

Per Hoffmann 34, 35, 47

Georg Homuth 56

Carsten Horn 57

Jouke-Jan Hottenga 10

David M Hougaard 9, 18

Marcus Ising 58

Rick Jansen 19

Ian Jones 59

Lisa A Jones 60

Eric Jorgenson 61

James A Knowles 62

Isaac S Kohane 63, 64, 65

Julia Kraft 4

Warren W. Kretzschmar 66

Jesper Krogh 67

Zoltán Kutalik 68, 69

Yihan Li 66

Penelope A Lind 28

Donald J MacIntyre 70, 71

Dean F MacKinnon 49

Robert M Maier 2

Wolfgang Maier 72

Jonathan Marchini 73

Hamdi Mbarek 10

Patrick McGrath 74

Peter McGuffin 27

Sarah E Medland 28

Divya Mehta 2, 75

Christel M Middeldorp 10, 76, 77

Evelin Mihailov 78

Yuri Milaneschi 19

Lili Milani 78

Francis M Mondimore 49

Grant W Montgomery 1

Sara Mostafavi 79, 80

Niamh Mullins 27

Matthias Nauck 81, 82

Bernard Ng 80

Michel G Nivard 10

Dale R Nyholt 83

Paul F O'Reilly 27

Hogni Oskarsson 84

Michael J Owen 59

Jodie N Painter 28

Carsten Bøcker Pedersen 9, 12, 13

Marianne Giørtz Pedersen 9, 12, 13

Roseann E. Peterson 17, 85

Erik Pettersson 22

Wouter J Peyrot 19

Giorgio Pistis 26

Danielle Posthuma 86, 87

Jorge A Quiroz 88

Per Qvist 7, 8, 9

John P Rice 89

Brien P. Riley 17

Margarita Rivera 27, 90

Saira Saeed Mirza 36

Robert Schoevers 91

Eva C Schulte 92, 93

Ling Shen 61

Jianxin Shi 94

Stanley I Shyn 95

Engilbert Sigurdsson 96

Grant C B Sinnamon 97

Johannes H Smit 19

Daniel J Smith 98

Hreinn Stefansson 99

Stacy Steinberg 99

Fabian Streit 45

Jana Strohmaier 45

Katherine E Tansey 100

Henning Teismann 101

Alexander Teumer 102

Wesley Thompson 9, 53, 103, 104

Pippa A Thomson 105

Thorgeir E Thorgeirsson 99

Matthew Traylor 106

Jens Treutlein 45

Vassily Trubetskoy 4

André G Uitterlinden 107

Daniel Umbricht 108

Sandra Van der Auwera 109

Albert M van Hemert 110

Alexander Viktorin 22

Peter M Visscher 1, 2

Yunpeng Wang 9, 53, 104

Bradley T. Webb 111

Shantel Marie Weinsheimer 9, 53

Jürgen Wellmann 101

Gonneke Willemsen 10

Stephanie H Witt 45

Yang Wu 1

Hualin S Xi 112

Jian Yang 2, 113

Futao Zhang 1

Volker Arolt 114

Bernhard T Baune 115

Klaus Berger 101

Dorret I Boomsma 10

Sven Cichon 34, 47, 116, 117

Udo Dannlowski 114

EJC de Geus 10, 118

J Raymond DePaulo 49

Enrico Domenici 119

Katharina Domschke 120

Tõnu Esko 5, 78

Hans J Grabe 109

Steven P Hamilton 121

Caroline Hayward 122

Andrew C Heath 89

Kenneth S Kendler 17

Stefan Kloiber 58, 123, 124

Glyn Lewis 125

Qingqin S Li 126

Susanne Lucae 58

Pamela AF Madden 89

Patrik K Magnusson 22

Nicholas G Martin 28

Andrew M McIntosh 11, 33

Andres Metspalu 78, 127

Ole Mors 9, 128

Preben Bo Mortensen 8, 9, 12, 13

Bertram Müller-Myhsok 15, 16, 129

Merete Nordentoft 9, 130

Markus M Nöthen 34, 35

Michael C O'Donovan 59

Sara A Paciga 131

Nancy L Pedersen 22

Brenda WJH Penninx 19

Roy H Perlis 38, 132

David J Porteous 105

James B Potash 133

Martin Preisig 26

Marcella Rietschel 45

Catherine Schaefer 61

Thomas G Schulze 45, 93, 134, 135, 136

Jordan W Smoller 38, 39, 40

Kari Stefansson 99, 137

Henning Tiemeier 36, 138, 139

Rudolf Uher 140

Henry Völzke 102

Myrna M Weissman 74, 141

Thomas Werge 9, 53, 142

Cathryn M Lewis 27, 143

Douglas F Levinson 144

Gerome Breen 27, 145

Anders D Børglum 7, 8, 9

Patrick F Sullivan 22, 146, 147,

1, Institute for Molecular Bioscience, The University of Queensland, Brisbane, QLD, AU

2, Queensland Brain Institute, The University of Queensland, Brisbane, QLD, AU

3, Analytic and Translational Genetics Unit, Massachusetts General Hospital, Boston, MA, US

4, Department of Psychiatry and Psychotherapy, Universitätsmedizin Berlin Campus Charité Mitte, Berlin, DE

5, Medical and Population Genetics, Broad Institute, Cambridge, MA, US

6, Centre for Psychiatry Research, Department of Clinical Neuroscience, Karolinska Institutet, Stockholm, SE

7, Department of Biomedicine, Aarhus University, Aarhus, DK

8, iSEQ, Centre for Integrative Sequencing, Aarhus University, Aarhus, DK

9, iPSYCH, The Lundbeck Foundation Initiative for Integrative Psychiatric Research,, DK

10, Dept of Biological Psychology & EMGO+ Institute for Health and Care Research, Vrije Universiteit Amsterdam, Amsterdam, NL

11, Division of Psychiatry, University of Edinburgh, Edinburgh, GB

12, Centre for Integrated Register-based Research, Aarhus University, Aarhus, DK

13, National Centre for Register-Based Research, Aarhus University, Aarhus, DK

14, Discipline of Psychiatry, University of Adelaide, Adelaide, SA, AU

15, Department of Translational Research in Psychiatry, Max Planck Institute of Psychiatry, Munich, DE

16, Munich Cluster for Systems Neurology (SyNergy), Munich, DE

17, Department of Psychiatry, Virginia Commonwealth University, Richmond, VA, US

18, Center for Neonatal Screening, Department for Congenital Disorders, Statens Serum Institut, Copenhagen, DK

19, Department of Psychiatry, Vrije Universiteit Medical Center and GGZ inGeest, Amsterdam, NL

20, Virginia Institute for Psychiatric and Behavior Genetics, Richmond, VA, US

21, Department of Psychiatry and Behavioral Sciences, Emory University School of Medicine, Atlanta, GA, US

22, Department of Medical Epidemiology and Biostatistics, Karolinska Institutet, Stockholm, SE

23, Department of Clinical Medicine, Translational Neuropsychiatry Unit, Aarhus University, Aarhus, DK

24, Human Genetics, Wellcome Trust Sanger Institute, Cambridge, GB

25, Statistical genomics and systems genetics, European Bioinformatics Institute (EMBL-EBI), Cambridge, GB

26, Department of Psychiatry, University Hospital of Lausanne, Prilly, Vaud, CH

27, Social Genetic and Developmental Psychiatry Centre, King's College London, London, GB

28, Genetics and Computational Biology, QIMR Berghofer Medical Research Institute, Brisbane, QLD, AU

29, Centre for Advanced Imaging, The University of Queensland, Brisbane, QLD, AU

30, Psychological Medicine, Cardiff University, Cardiff, GB

31, Center for Genomic and Computational Biology, Duke University, Durham, NC, US

32, Department of Pediatrics, Division of Medical Genetics, Duke University, Durham, NC, US

33, Centre for Cognitive Ageing and Cognitive Epidemiology, University of Edinburgh, Edinburgh, GB

34, Institute of Human Genetics, University of Bonn, Bonn, DE

35, Life&Brain Center, Department of Genomics, University of Bonn, Bonn, DE

36, Epidemiology, Erasmus MC, Rotterdam, Zuid-Holland, NL

37, Psychiatry, Dokuz Eylul University School Of Medicine, Izmir, TR

38, Department of Psychiatry, Massachusetts General Hospital, Boston, MA, US

39, Psychiatric and Neurodevelopmental Genetics Unit (PNGU), Massachusetts General Hospital, Boston, MA, US

40, Stanley Center for Psychiatric Research, Broad Institute, Cambridge, MA, US

41, Neuroscience and Mental Health, Cardiff University, Cardiff, GB

42, Bioinformatics, University of British Columbia, Vancouver, BC, CA

43, Department of Epidemiology, Harvard T.H. Chan School of Public Health, Boston, MA, US

44, Department of Mathematics, Massachusetts Institute of Technology, Cambridge, MA, US

45, Department of Genetic Epidemiology in Psychiatry, Central Institute of Mental Health,  Medical Faculty Mannheim, Heidelberg University, Mannheim, Baden-Württemberg, DE

46, Department of Psychiatry (UPK), University of Basel, Basel, CH

47, Human Genomics Research Group, Department of Biomedicine, University of Basel, Basel, CH

48, Department of Psychiatry, Trinity College Dublin, Dublin, IE

49, Psychiatry & Behavioral Sciences, Johns Hopkins University, Baltimore, MD, US

50, Bioinformatics Research Centre, Aarhus University, Aarhus, DK

51, Institute of Genetic Medicine, Newcastle University, Newcastle upon Tyne, GB

52, Danish Headache Centre, Department of Neurology, Rigshospitalet, Glostrup, DK

53, Institute of Biological Psychiatry, Mental Health Center Sct. Hans, Mental Health Services Capital Region of Denmark, Copenhagen, DK

54, iPSYCH, The Lundbeck Foundation Initiative for Psychiatric Research, Copenhagen, DK

55, Brain and Mind Centre, University of Sydney, Sydney, NSW, AU

56, Interfaculty Institute for Genetics and Functional Genomics, Department of Functional Genomics, University Medicine and Ernst Moritz Arndt University Greifswald, Greifswald, Mecklenburg-Vorpommern, DE

57, Roche Pharmaceutical Research and Early Development, Pharmaceutical Sciences, Roche Innovation Center Basel, F. Hoffmann-La Roche Ltd, Basel, CH

58, Max Planck Institute of Psychiatry, Munich, DE

59, MRC Centre for Neuropsychiatric Genetics and Genomics, Cardiff University, Cardiff, GB

60, Department of Psychological Medicine, University of Worcester, Worcester, GB

61, Division of Research, Kaiser Permanente Northern California, Oakland, CA, US

62, Psychiatry & The Behavioral Sciences, University of Southern California, Los Angeles, CA, US

63, Department of Biomedical Informatics, Harvard Medical School, Boston, MA, US

64, Department of Medicine, Brigham and Women's Hospital, Boston, MA, US

65, Informatics Program, Boston Children's Hospital, Boston, MA, US

66, Wellcome Trust Centre for Human Genetics, University of Oxford, Oxford, GB

67, Department of Endocrinology at Herlev University Hospital, University of Copenhagen, Copenhagen, DK

68, Institute of Social and Preventive Medicine (IUMSP), University Hospital of Lausanne, Lausanne, VD, CH

69, Swiss Institute of Bioinformatics, Lausanne, VD, CH

70, Division of Psychiatry, Centre for Clinical Brain Sciences, University of Edinburgh, Edinburgh, GB

71, Mental Health, NHS 24, Glasgow, GB

72, Department of Psychiatry and Psychotherapy, University of Bonn, Bonn, DE

73, Statistics, University of Oxford, Oxford, GB

74, Psychiatry, Columbia University College of Physicians and Surgeons, New York, NY, US

75, School of Psychology and Counseling, Queensland University of Technology, Brisbane, QLD, AU

76, Child and Youth Mental Health Service, Children's Health Queensland Hospital and Health Service, South Brisbane, QLD, AU

77, Child Health Research Centre, University of Queensland, Brisbane, QLD, AU

78, Estonian Genome Center, University of Tartu, Tartu, EE

79, Medical Genetics, University of British Columbia, Vancouver, BC, CA

80, Statistics, University of British Columbia, Vancouver, BC, CA

81, DZHK (German Centre for Cardiovascular Research), Partner Site Greifswald, University Medicine, University Medicine Greifswald, Greifswald, Mecklenburg-Vorpommern, DE

82, Institute of Clinical Chemistry and Laboratory Medicine, University Medicine Greifswald, Greifswald, Mecklenburg-Vorpommern, DE

83, Institute of Health and Biomedical Innovation, Queensland University of Technology, Brisbane, QLD, AU

84, Humus, Reykjavik, IS

85, Virginia Institute for Psychiatric & Behavioral Genetics, Virginia Commonwealth University, Richmond, VA, US

86, Clinical Genetics, Vrije Universiteit Medical Center, Amsterdam, NL

87, Complex Trait Genetics, Vrije Universiteit Amsterdam, Amsterdam, NL

88, Solid Biosciences, Boston, MA, US

89, Department of Psychiatry, Washington University in Saint Louis School of Medicine, Saint Louis, MO, US

90, Department of Biochemistry and Molecular Biology II, Institute of Neurosciences, Center for Biomedical Research, University of Granada, Granada, ES

91, Department of Psychiatry, University of Groningen, University Medical Center Groningen, Groningen, NL

92, Department of Psychiatry and Psychotherapy, Medical Center of the University of Munich, Campus Innenstadt, Munich, DE

93, Institute of Psychiatric Phenomics and Genomics (IPPG), Medical Center of the University of Munich, Campus Innenstadt, Munich, DE

94, Division of Cancer Epidemiology and Genetics, National Cancer Institute, Bethesda, MD, US

95, Behavioral Health Services, Kaiser Permanente Washington, Seattle, WA, US

96, Faculty of Medicine, Department of Psychiatry, University of Iceland, Reykjavik, IS

97, School of Medicine and Dentistry, James Cook University, Townsville, QLD, AU

98, Institute of Health and Wellbeing, University of Glasgow, Glasgow, GB

99, deCODE Genetics / Amgen, Reykjavik, IS

100, College of Biomedical and Life Sciences, Cardiff University, Cardiff, GB

101, Institute of Epidemiology and Social Medicine, University of Münster, Münster, Nordrhein-Westfalen, DE

102, Institute for Community Medicine, University Medicine Greifswald, Greifswald, Mecklenburg-Vorpommern, DE

103, Department of Psychiatry, University of California, San Diego, San Diego, CA, US

104, KG Jebsen Centre for Psychosis Research, Norway Division of Mental Health and Addiction, Oslo University Hospital, Oslo, NO

105, Medical Genetics Section, CGEM, IGMM, University of Edinburgh, Edinburgh, GB

106, Clinical Neurosciences, University of Cambridge, Cambridge, GB

107, Internal Medicine, Erasmus MC, Rotterdam, Zuid-Holland, NL

108, Roche Pharmaceutical Research and Early Development, Neuroscience, Ophthalmology and Rare Diseases Discovery & Translational Medicine Area, Roche Innovation Center Basel, F. Hoffmann-La Roche Ltd, Basel, CH

109, Department of Psychiatry and Psychotherapy, University Medicine Greifswald, Greifswald, Mecklenburg-Vorpommern, DE

110, Department of Psychiatry, Leiden University Medical Center, Leiden, NL

111, Virginia Institute for Psychiatric & Behavioral Genetics, Virginia Commonwealth University, Richmond, VA, US

112, Computational Sciences Center of Emphasis, Pfizer Global Research and Development, Cambridge, MA, US

113, Institute for Molecular Bioscience; Queensland Brain Institute, The University of Queensland, Brisbane, QLD, AU

114, Department of Psychiatry, University of Münster, Münster, Nordrhein-Westfalen, DE

115, Department of Psychiatry, Melbourne Medical School, University of Melbourne, Melbourne, AU

116, Institute of Medical Genetics and Pathology, University Hospital Basel, University of Basel, Basel, CH

117, Institute of Neuroscience and Medicine (INM-1), Research Center Juelich, Juelich, DE

118, Amsterdam Public Health Institute, Vrije Universiteit Medical Center, Amsterdam, NL

119, Centre for Integrative Biology, Università degli Studi di Trento, Trento, Trentino-Alto Adige, IT

120, Department of Psychiatry and Psychotherapy, Medical Center, University of Freiburg, Faculty of Medicine, University of Freiburg, Freiburg, DE

121, Psychiatry, Kaiser Permanente Northern California, San Francisco, CA, US

122, Medical Research Council Human Genetics Unit, Institute of Genetics and Molecular Medicine, University of Edinburgh, Edinburgh, GB

123, Department of Psychiatry, University of Toronto, Toronto, ON, CA

124, Centre for Addiction and Mental Health, Toronto, ON, CA

125, Division of Psychiatry, University College London, London, GB

126, Neuroscience Therapeutic Area, Janssen Research and Development, LLC, Titusville, NJ, US

127, Institute of Molecular and Cell Biology, University of Tartu, Tartu, EE

128, Psychosis Research Unit, Aarhus University Hospital, Risskov, Aarhus, DK

129, University of Liverpool, Liverpool, GB

130, Mental Health Center Copenhagen, Copenhagen Universtity Hospital, Copenhagen, DK

131, Human Genetics and Computational Biomedicine, Pfizer Global Research and Development, Groton, CT, US

132, Psychiatry, Harvard Medical School, Boston, MA, US

133, Psychiatry, University of Iowa, Iowa City, IA, US

134, Department of Psychiatry and Behavioral Sciences, Johns Hopkins University, Baltimore, MD, US

135, Department of Psychiatry and Psychotherapy, University Medical Center Göttingen, Goettingen, Niedersachsen, DE

136, Human Genetics Branch, NIMH Division of Intramural Research Programs, Bethesda, MD, US

137, Faculty of Medicine, University of Iceland, Reykjavik, IS

138, Child and Adolescent Psychiatry, Erasmus MC, Rotterdam, Zuid-Holland, NL

139, Psychiatry, Erasmus MC, Rotterdam, Zuid-Holland, NL

140, Psychiatry, Dalhousie University, Halifax, NS, CA

141, Division of Epidemiology, New York State Psychiatric Institute, New York, NY, US

142, Department of Clinical Medicine, University of Copenhagen, Copenhagen, DK

143, Department of Medical & Molecular Genetics, King's College London, London, GB

144, Psychiatry & Behavioral Sciences, Stanford University, Stanford, CA, US

145, NIHR Maudsley Biomedical Research Centre, King's College London, London, GB

146, Genetics, University of North Carolina at Chapel Hill, Chapel Hill, NC, US

147, Psychiatry, University of North Carolina at Chapel Hill, Chapel Hill, NC, US

Version 5. 2018-07-17

### Acknowledgments

The 1^st^ author AAS is funded by University of Edinburgh ([www.ed.ac.uk](http://www.ed.ac.uk)) and Medical Research Council for his PhD study at the University of Edinburgh Institute of Genetics and Molecular Medicine ([www.ed.ac.uk/igmm](http://www.ed.ac.uk/igmm)). MA is supported by STRADL through a Wellcome Trust Strategic Award (reference 104036/Z/14/Z). DJM acknowledges the financial support of NHS Research Scotland (NRS), through NHS Lothian. Generation Scotland received core support from the Chief Scientist Office of the Scottish Government Health Directorates [CZD/16/6] and the Scottish Funding Council [HR03006]. Genotyping of the Generation Scotland samples was carried out by the Genetics Core Laboratory at the Wellcome Trust Clinical Research Facility, Edinburgh, Scotland and was funded by the Medical Research Council UK and the Wellcome Trust (Wellcome Trust Strategic Award “STratifying Resilience and Depression Longitudinally” (STRADL), Reference 104036/Z/14/Z). The Psychiatric Genomics Consortium has received major funding from the US National Institute of Mental Health and the US National Institute of Drug Abuse (U01 MH109528 and U01 MH1095320).

### Supplementary Figures

#### Supplementary Figure 1. Manhattan and QQ plots of Generation Scotland GWAS and GWEIS using TSLE as exposure

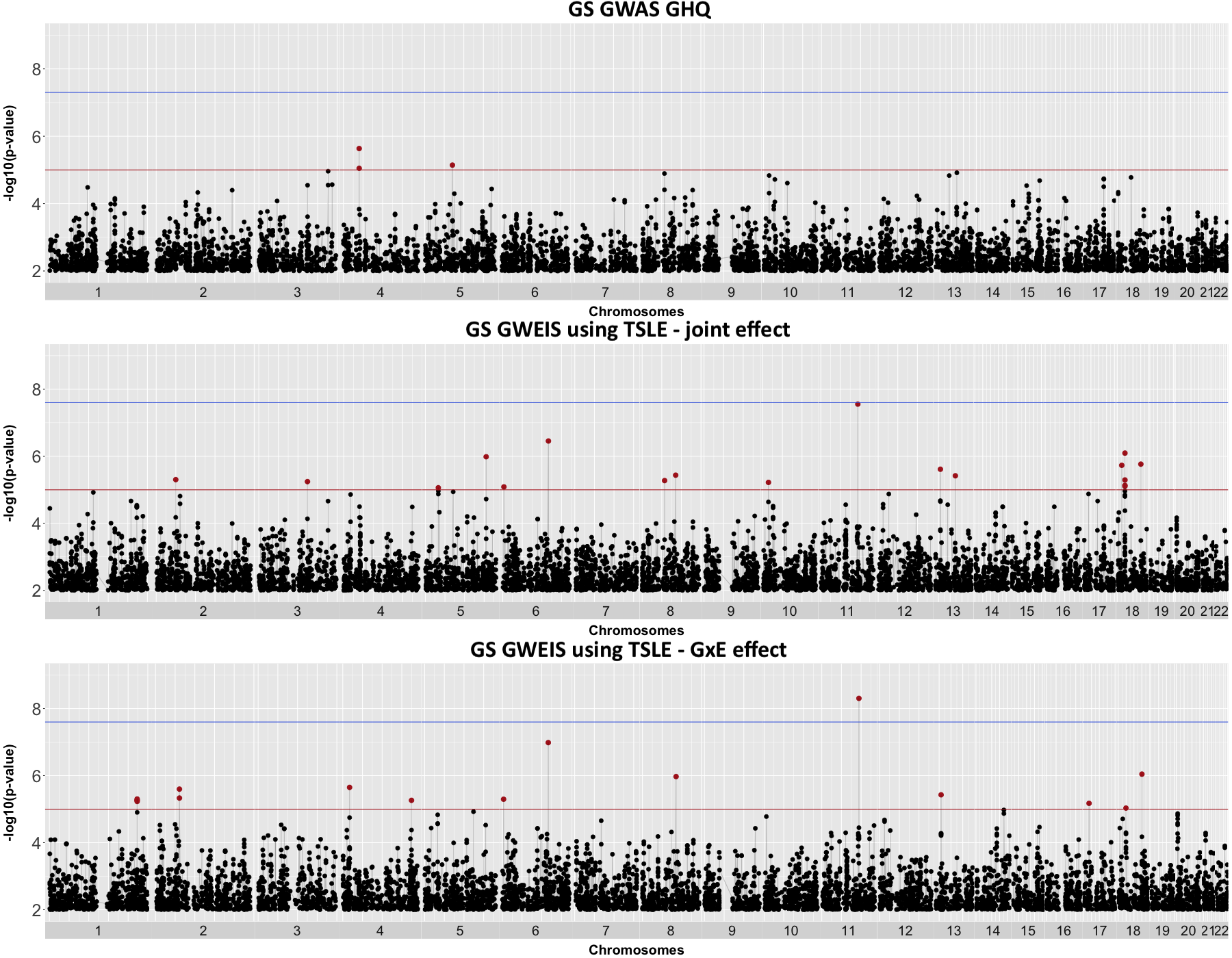

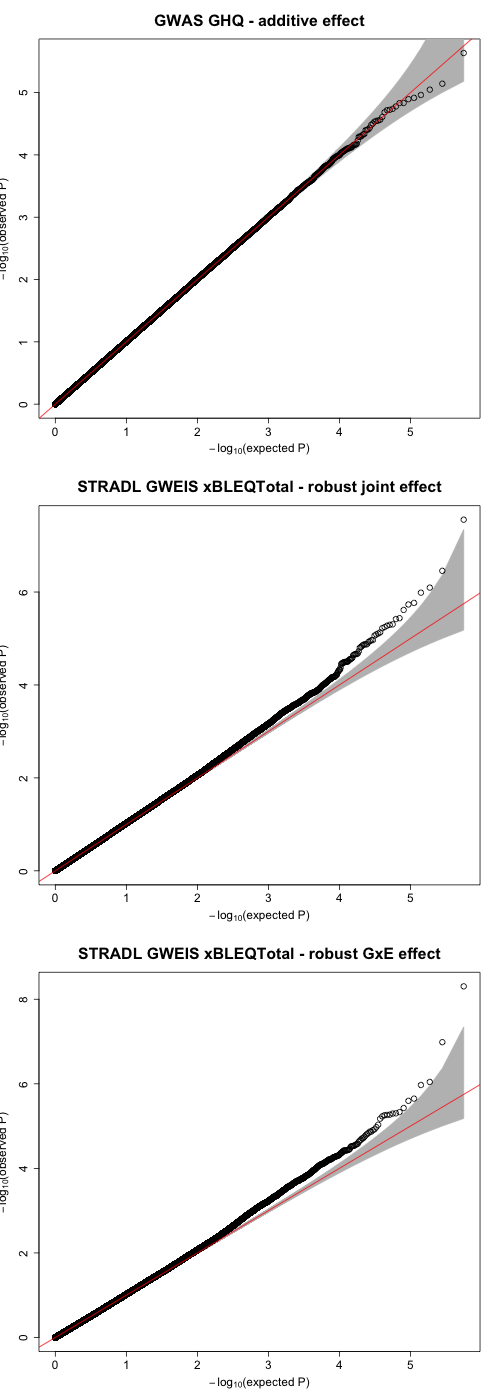

**λ_1000_=1.007972**

λ=1.039213

SE=3.156605e^-5^

**λ_1000_=1.007448**

λ=1.036637

SE=2.084289e^-5^

**λ_1000_=1.000879**

λ=1.004322

SE=4.624774e^-6^

Manhattan plots showing associations of GHQ in Generation Scotland (N = 4,919) from (top) additive effects from GWAS, (middle) joint effect from GWEIS and (bottom) GxE effect from GWEIS, using “total” SLE (TSLE) as exposure. Suggestive genome-wide significance threshold (*p =* 1x10^-5^) is shown by solid red line. Genome-wide significance threshold (GWAS: *p* = 5x10^-8^; GWEIS: *p* = 2.97x10^-8^) is shown by solid blue line. QQ plots on the right show genomic factor inflation and represented by lambda estimates λ and λ_1000_; grey shadow represents the 95% confidence intervals.

#### Supplementary Figure 2. Manhattan and QQ plots of Generation Scotland GWAS and GWEIS using DSLE as exposure

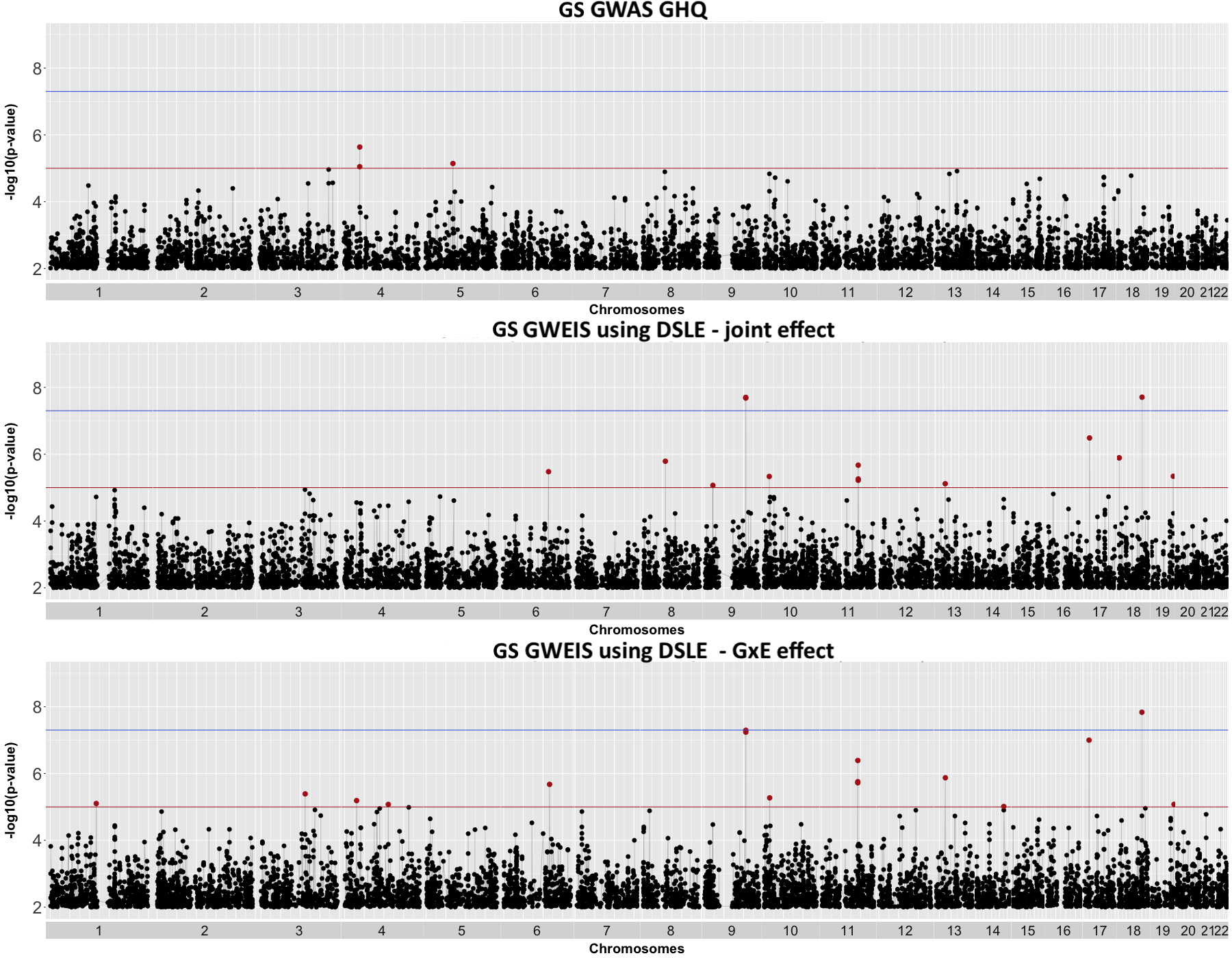

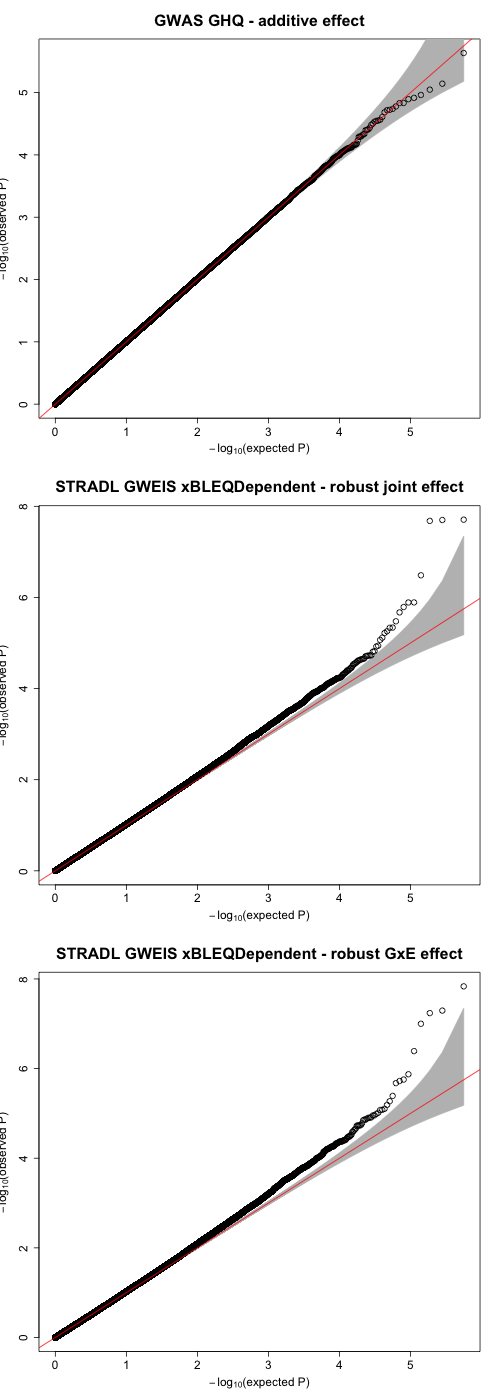

**λ_1000_=1.008821**

λ=1.043391

SE=3.185847e^-5^

**λ_1000_=1.008176**

λ=1.040219

SE=2.520509e^-5^

**λ_1000_=1.000879**

λ=1.004322

SE=4.624774e^-6^

Manhattan plots showing associations of GHQ in Generation Scotland (N = 4,919) from (top) additive effects from GWAS, (middle) joint effect from GWEIS and (bottom) GxE effect from GWEIS, using “dependent” SLE (DSLE) as exposure. Suggestive genome-wide significance threshold (*p =* 1x10^-5^) is shown by solid red line. Genome-wide significance threshold (GWAS: *p* = 5x10^-8^; GWEIS: *p* = 2.97x10^-8^) is shown by solid blue line. QQ plots on the right show genomic factor inflation and represented by lambda estimates λ and λ_1000_; grey shadow represents the 95% confidence intervals.

#### Supplementary Figure 3. Manhattan and QQ plots of Generation Scotland GWAS and GWEIS using ISLE as exposure

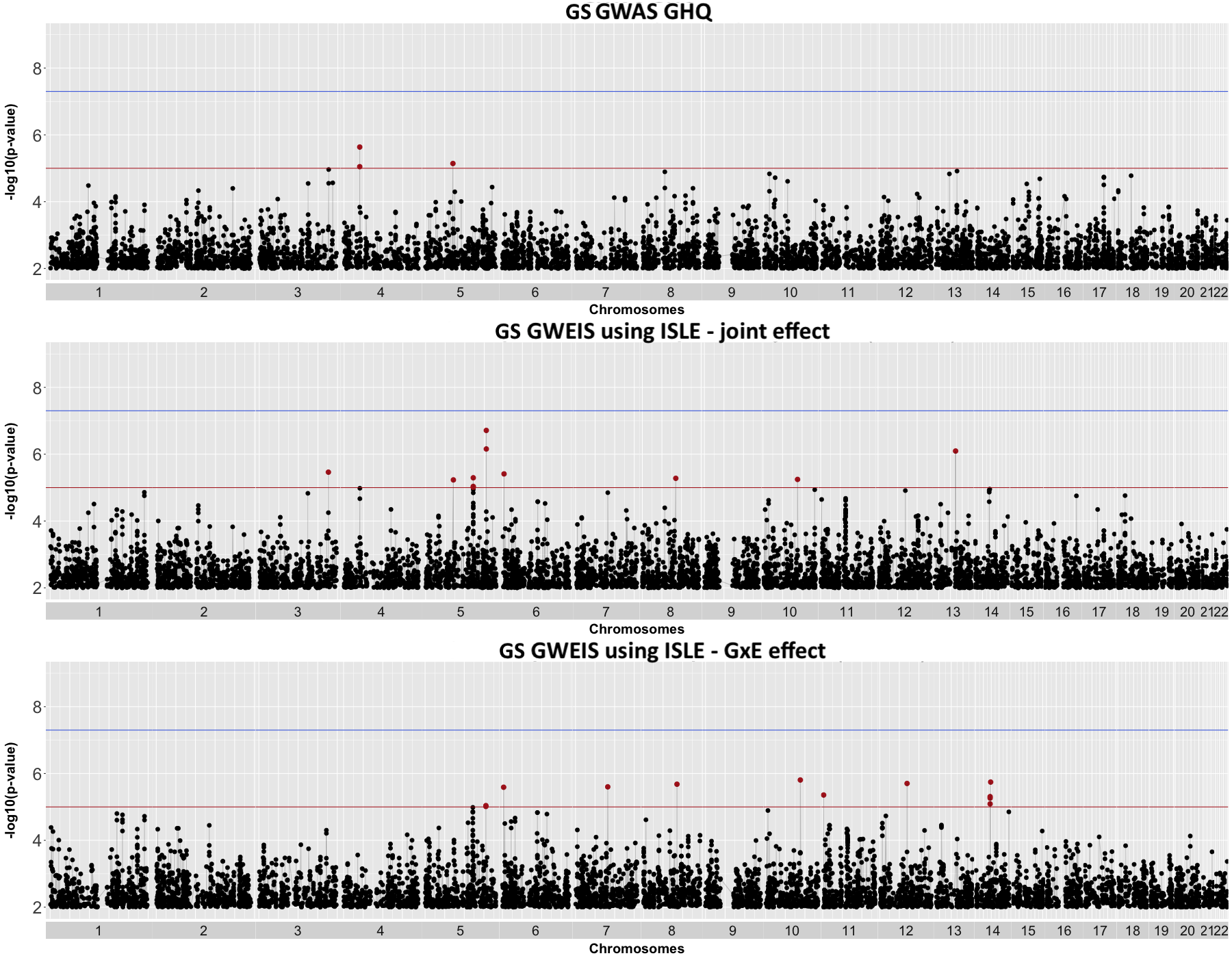

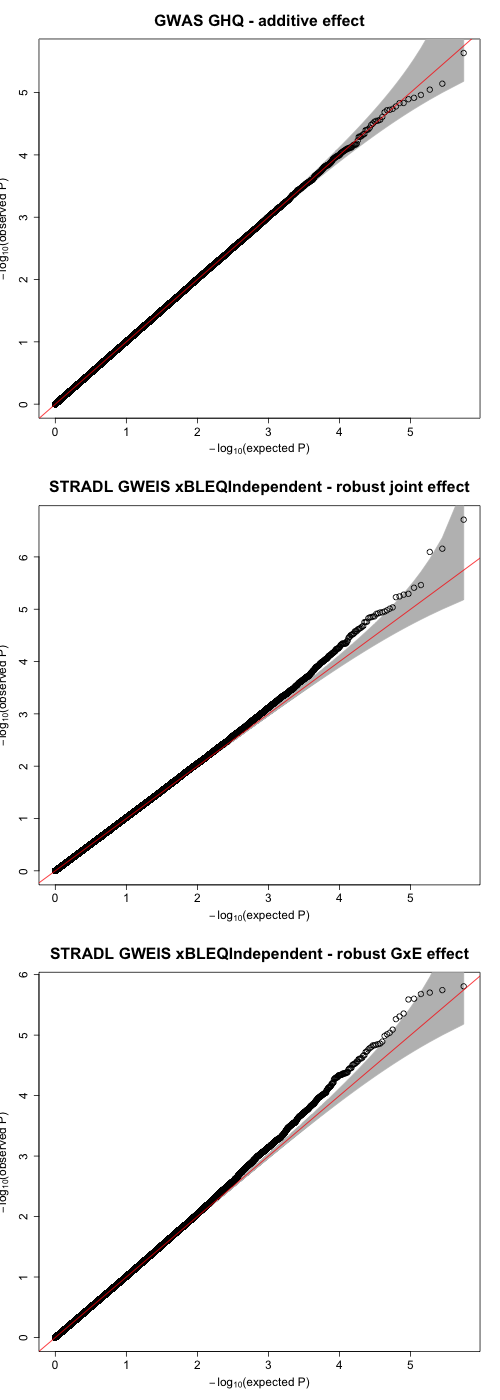

**λ_1000_=1.004788**

λ=1.023551

SE=2.17667e^-5^

**λ_1000_=1.005561**

λ=1.027353

SE=1.398843e^-5^

**λ_1000_=1.000879**

λ=1.004322

SE=4.624774e^-6^

Manhattan plots showing associations of GHQ in Generation Scotland (N = 4,919) from (top) additive effects from GWAS, (middle) joint effect from GWEIS and (bottom) GxE effect from GWEIS, using “independent” SLE (ISLE) as exposure. Suggestive genome-wide significance threshold (*p =* 1x10^-5^) is shown by solid red line. Genome-wide significance threshold (GWAS: *p* = 5x10^-8^; GWEIS: *p* = 2.97x10^-8^) is shown by solid blue line. QQ plots on the right show genomic factor inflation and represented by lambda estimates λ and λ_1000_; grey shadow represents 95% confidence intervals.

#### Supplementary Figure 4. Manhattan and QQ plots of UK Biobank GWAS and GWEIS

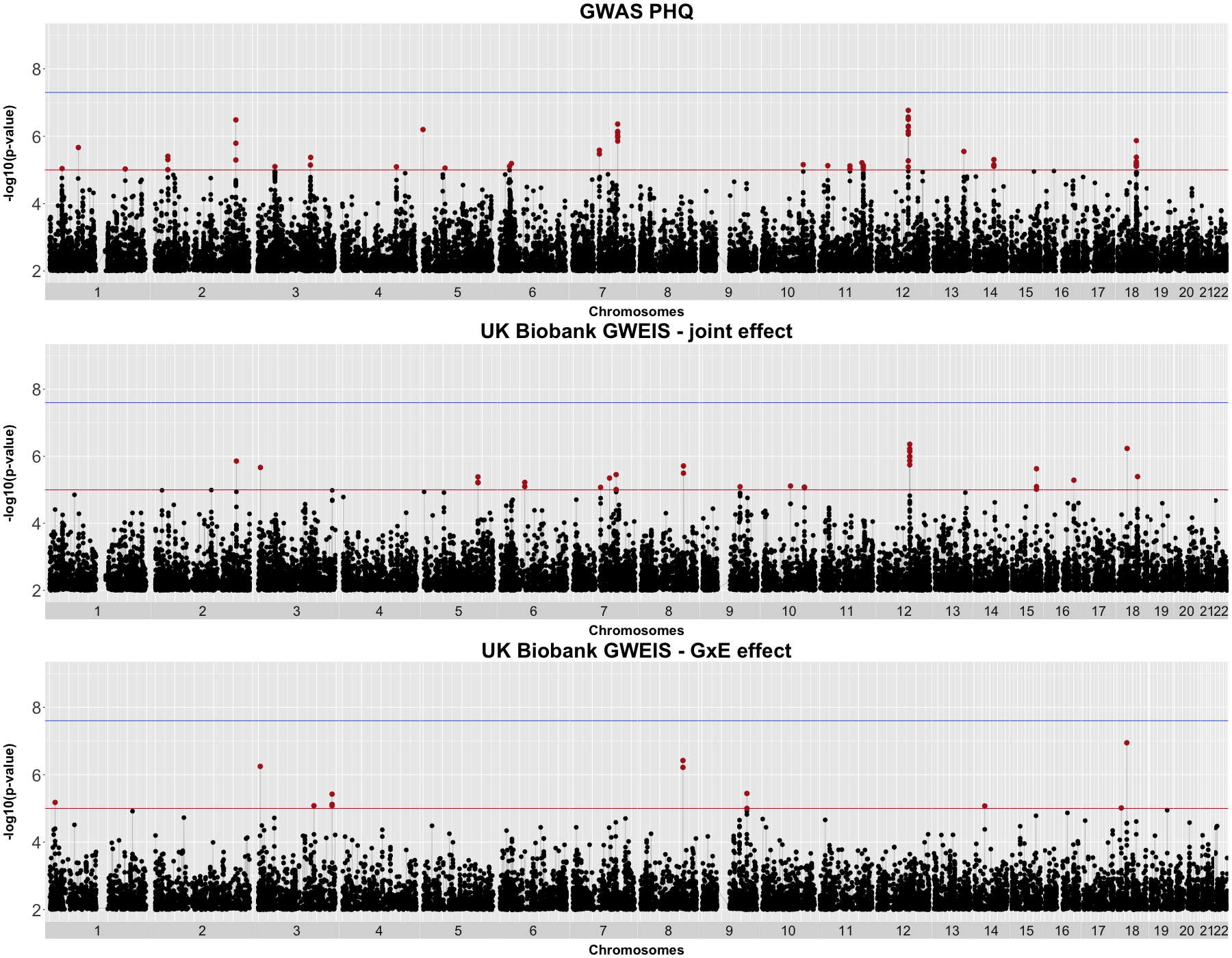

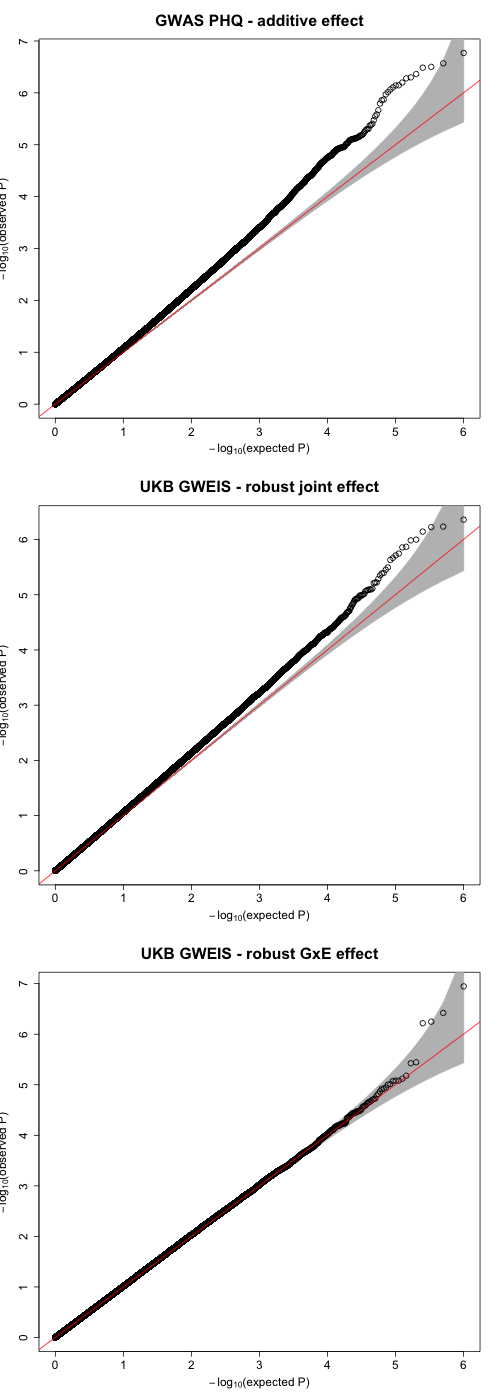

**λ_1000_=1.000156**

λ=1.015425

SE=4.964996e^-6^

**λ_1000_=1.000826**

λ=1.08186

SE=5.479599e^-6^

**λ_1000_=1.00123**

λ=1.121865

SE=2.201649e^-5^

Manhattan plots showing associations of PHQ in UK Biobank (N = 99,057) from (top) additive effects from GWAS, (middle) joint effect from GWEIS and (bottom) GxE effect from GWEIS, using “total” SLE as exposure. Suggestive genome-wide significance threshold (*p =* 1x10^-5^) is shown by solid red line. Genome-wide significance threshold (GWAS: *p* = 5x10^-8^; GWEIS: *p* = 2.47x10^-8^) is shown by solid blue line. QQ plots on the right show genomic factor inflation and represented by lambda estimates λ and λ_1000_; grey shadow represents the 95% confidence intervals.

#### Supplementary Figure 5. GxE effect derived from GWEIS on Generation Scotland using TSLE as exposure, regional plot of locus around rs12789145 on chromosome 11

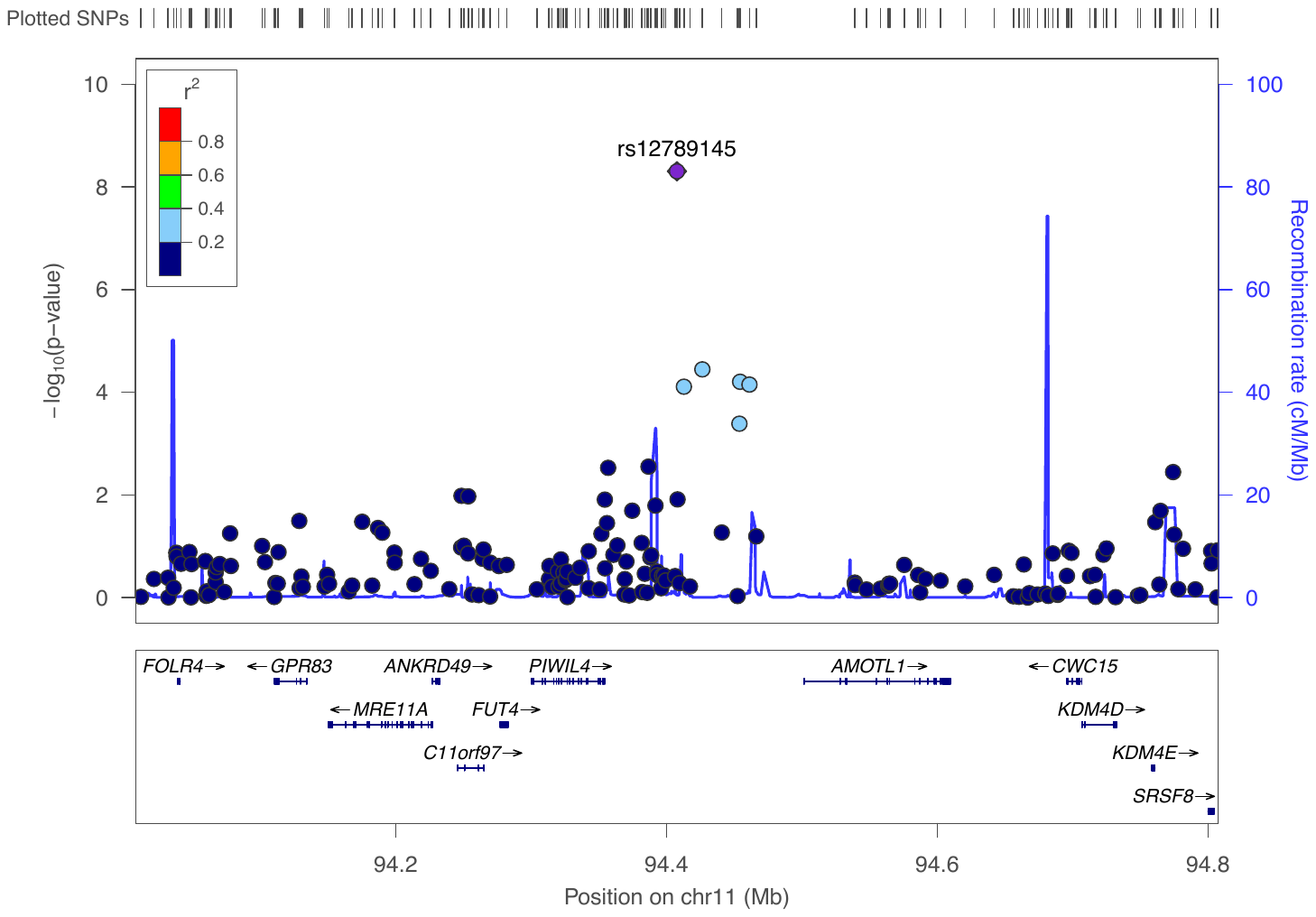

LocusZoom plot of the associated region on chromosome 11. Genotyped SNPs are shown as circles. Linkage disequilibrium structure across the region is represented by colour sclae (r^2^). Recombination rate is indicated by solid blue line. Y-axis shows significance association with GxE effect derived from GWEIS on Generation Scotland using TSLE as exposure. Location, structure and transcription direction of the genes in the region are represented below.

#### Supplementary Figure 6. GxE effect derived from GWEIS on Generation Scotland using DSLE as exposure, regional plot of locus around rs17070072 on chromosome 18

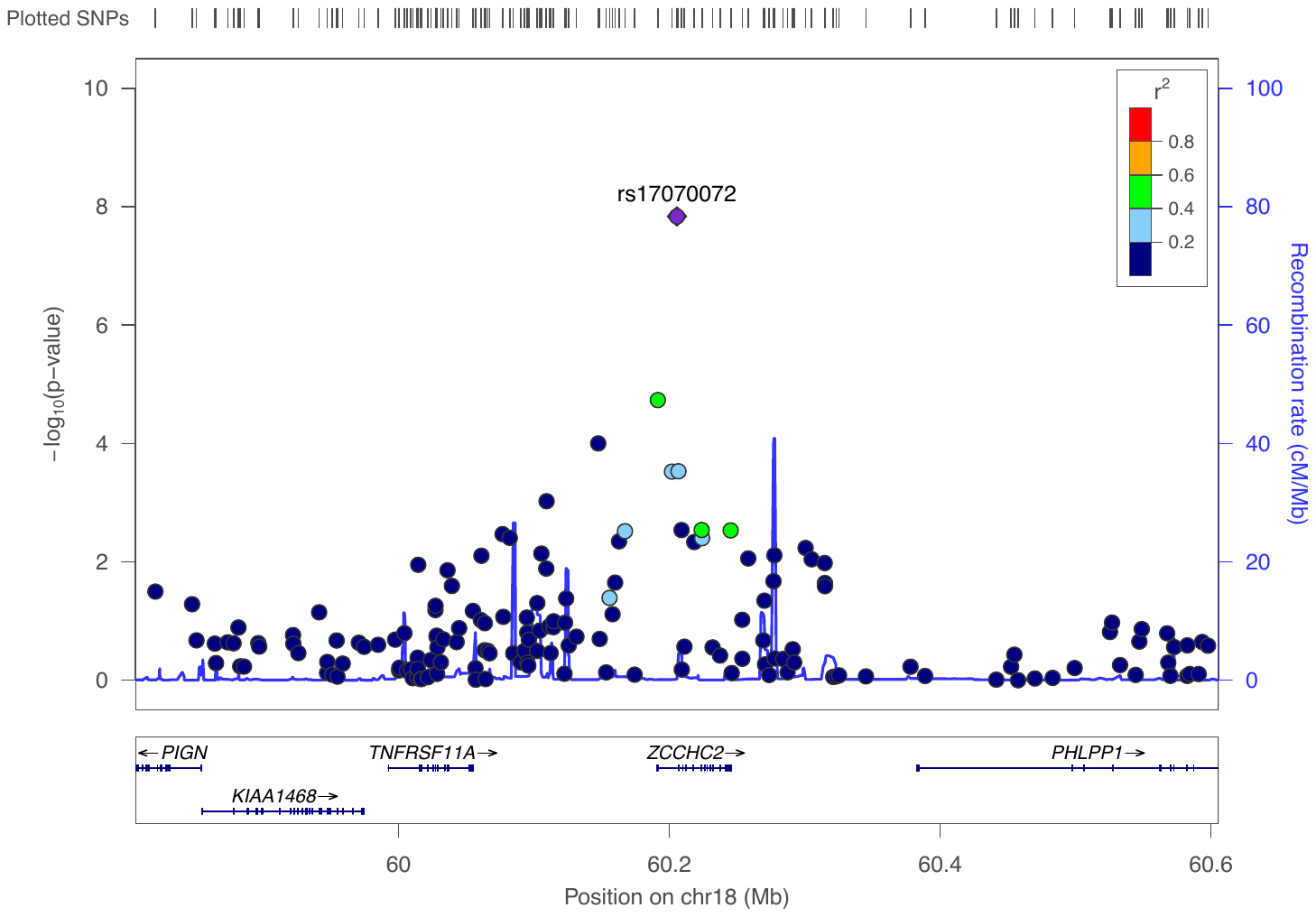

LocusZoom plot of the associated region on chromosome 18. Genotyped SNPs are shown as circles. Linkage disequilibrium structure across the region is represented by colour sclae (r^2^). Recombination rate is indicated by solid blue line. Y-axis shows significance association with GxE effect derived from GWEIS on Generation Scotland using DSLE as exposure. Location, structure and transcription direction of the genes in the region are represented below.

#### Supplementary Figure 7. Joint effect derived from GWEIS on Generation Scotland using DSLE as exposure, regional plot of locus around rs17070072 on chromosome 9

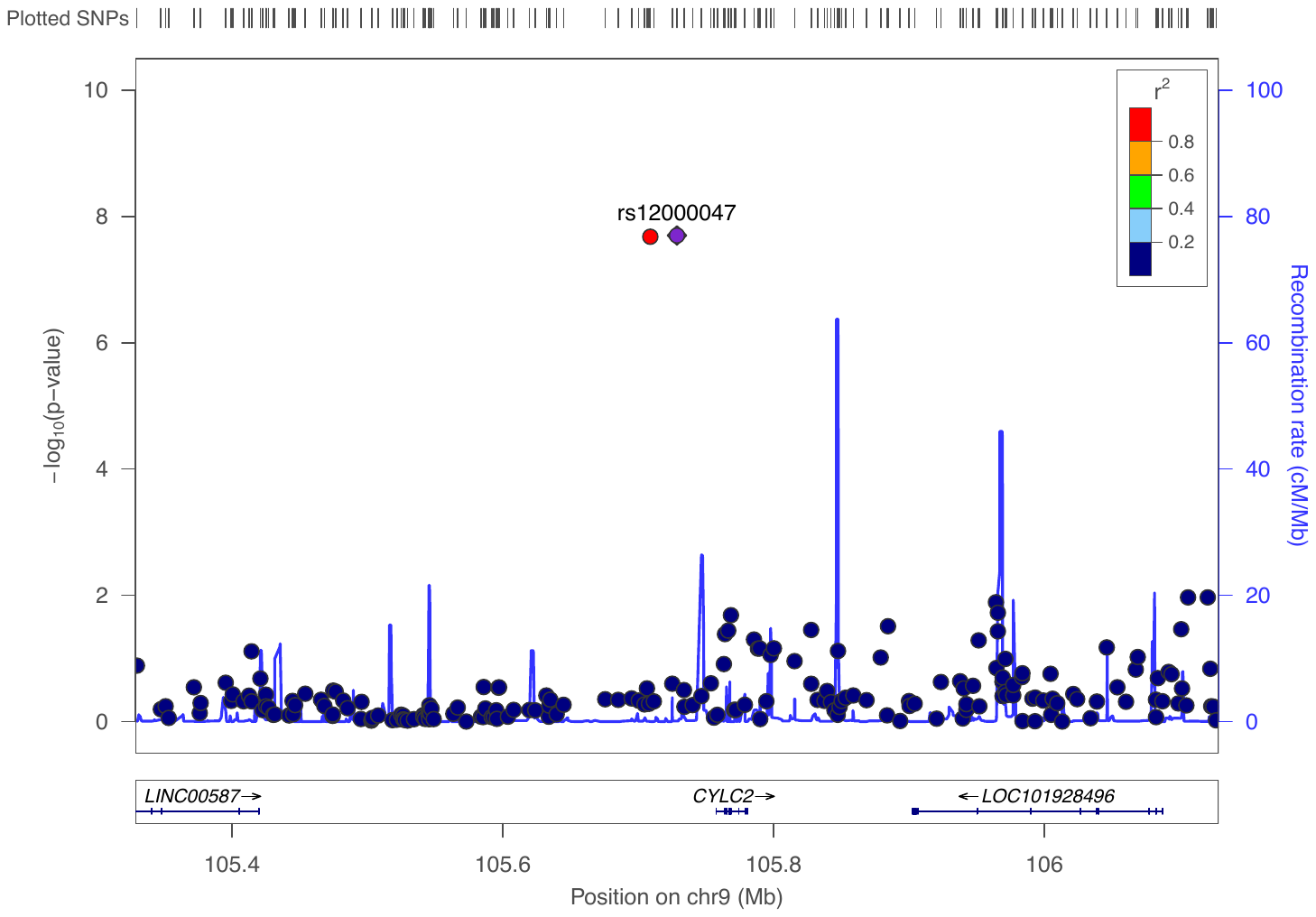

LocusZoom plot of the associated region on chromosome 9. Genotyped SNPs are shown as circles. Linkage disequilibrium structure across the region is represented by colour sclae (r^2^). Recombination rate is indicated by solid blue line. Y-axis shows significance association with joint effect derived from GWEIS on Generation Scotland using DSLE as exposure. Location, structure and transcription direction of the genes in the region are represented below.

#### Supplementary Figure 8. Manhattan plot of gene-based test of PHQ UKB GWAS

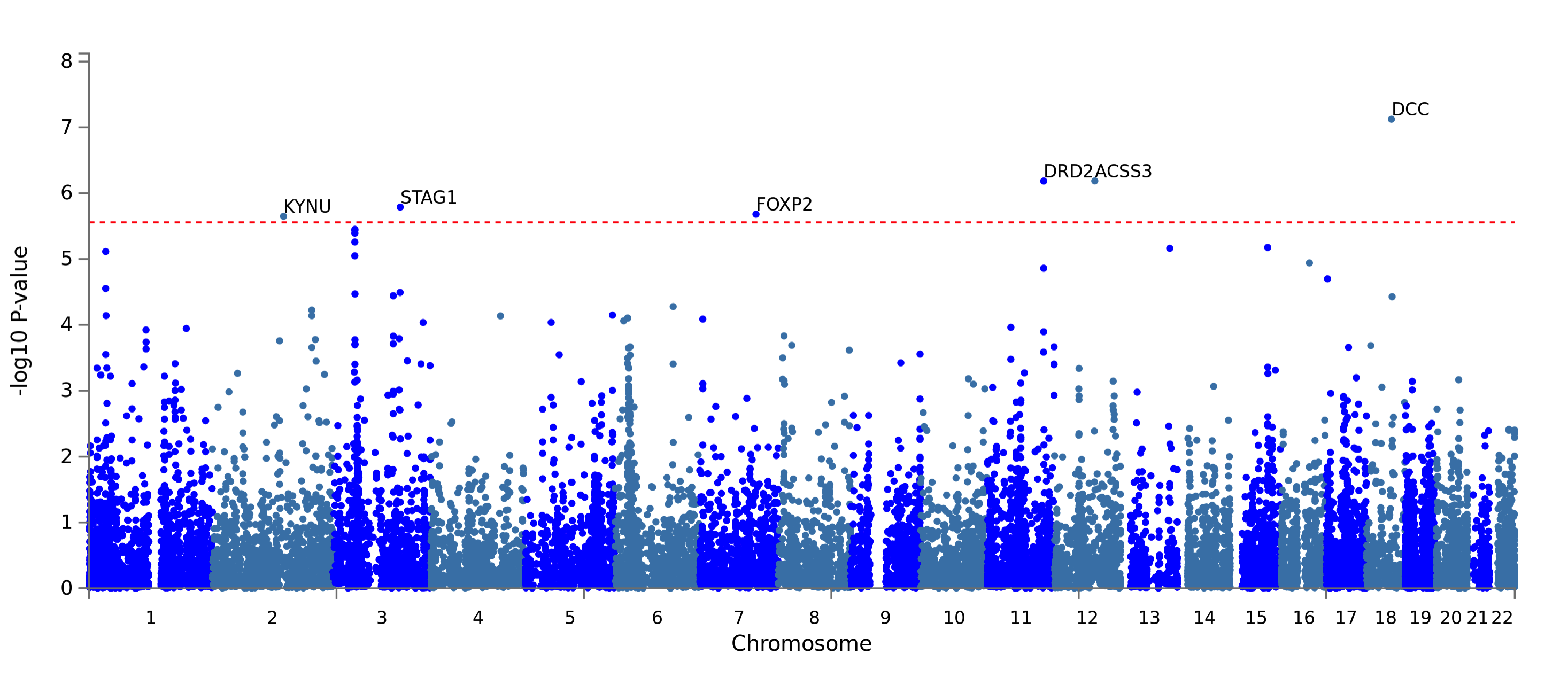

Manhattan plot showing gene-based association of PHQ using summary statistics for the additive effect derived from GWAS in UK Biobank (N = 99,057). The x-axis is base-paired chromosomal position and the y-axis represents the significance (-log_10_ *p* value) of association between main additive effects and PHQ. Bonferroni-corrected significance threshold (*p* = 0.05 / 18,068 = 2.77x10^-6^) is shown by dashed red line.

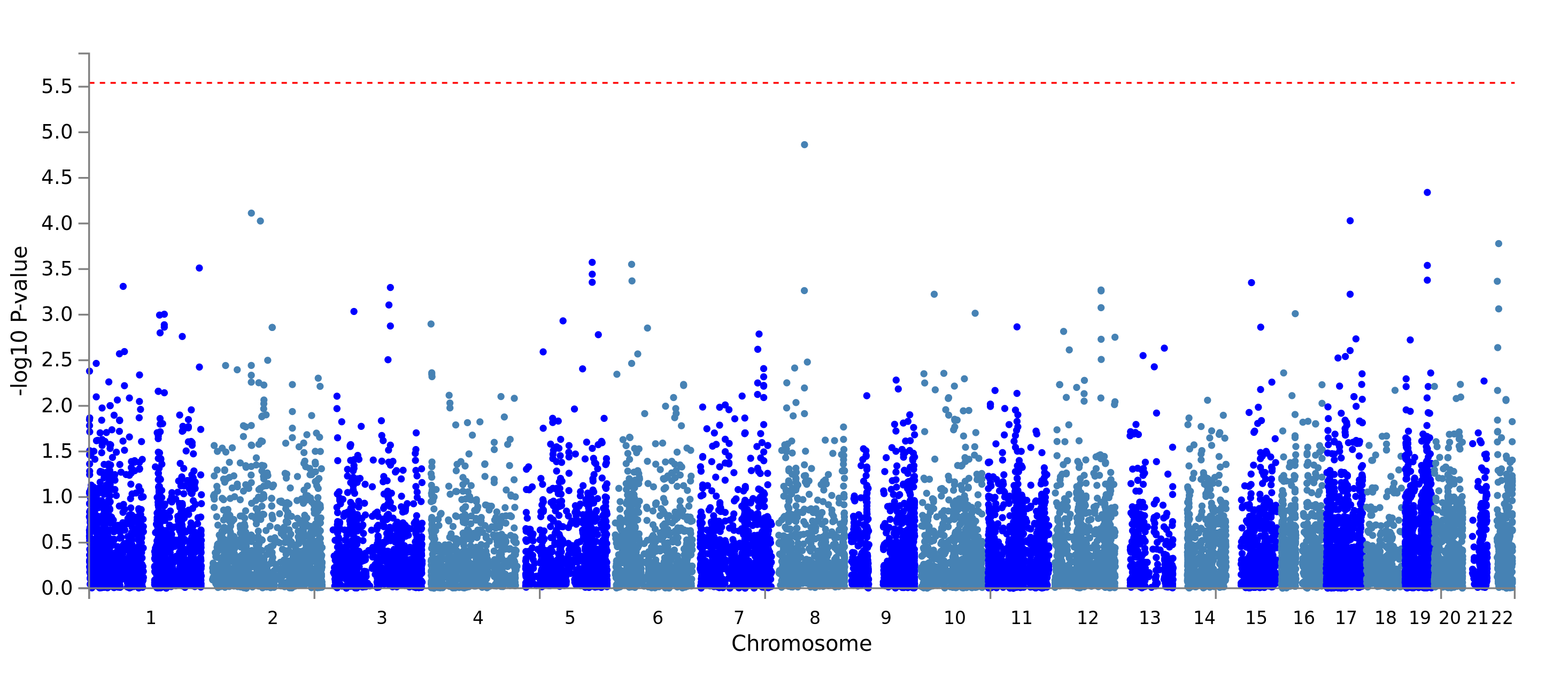

#### Supplementary Figure 9. Manhattan plot of gene-based test of GHQ GS GWAS

Manhattan plot showing gene-based association of GHQ using summary statistics for the additive effect derived from GWAS in Generation Scotland (N = 4,919). The x-axis is base-paired chromosomal position and the y-axis represents the significance (-log_10_ *p* value) of association between main additive effects and GHQ. Bonferroni-corrected significance threshold (*p* = 0.05 / 18,068 = 2.77x10^-6^) is shown by dashed red line.

#### Supplementary Figure 10. Manhattan plot of gene-based test of PHQ UKB GWEIS using TSLE_UKB_ as exposure for GxE effect

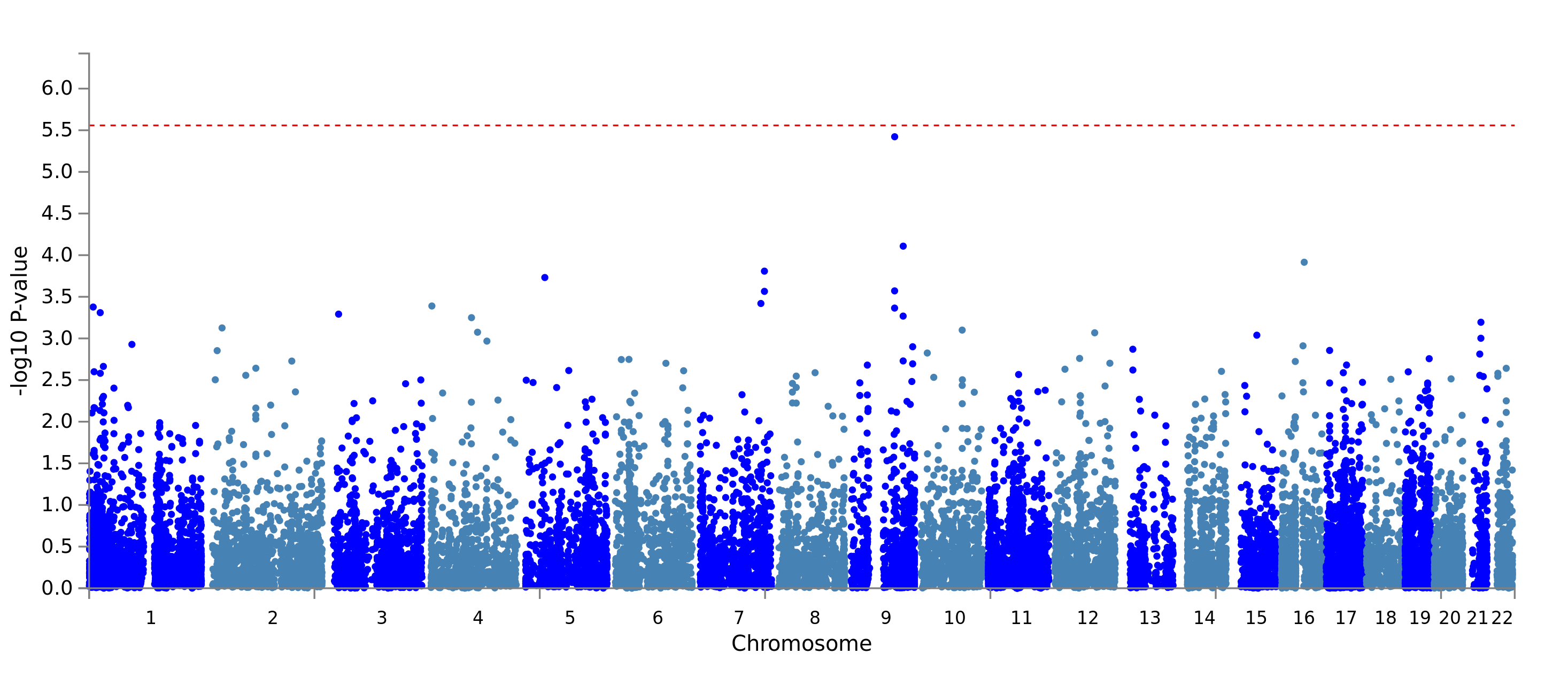

Manhattan plot showing gene-based association of PHQ using summary statistics for the GxE effect derived from GWEIS in UK Biobank (N = 99,057). The x-axis is base-paired chromosomal position and the y-axis represents the significance (-log_10_ *p* value) of association between GxE effects and PHQ. Bonferroni-corrected significance threshold (*p* = 0.05 / 18,068 = 2.77x10^-6^) is shown by dashed red line.

#### Supplementary Figure 11. Manhattan plot of gene-based test of PHQ UKB GWEIS using TSLE_UKB_ as exposure for joint effect

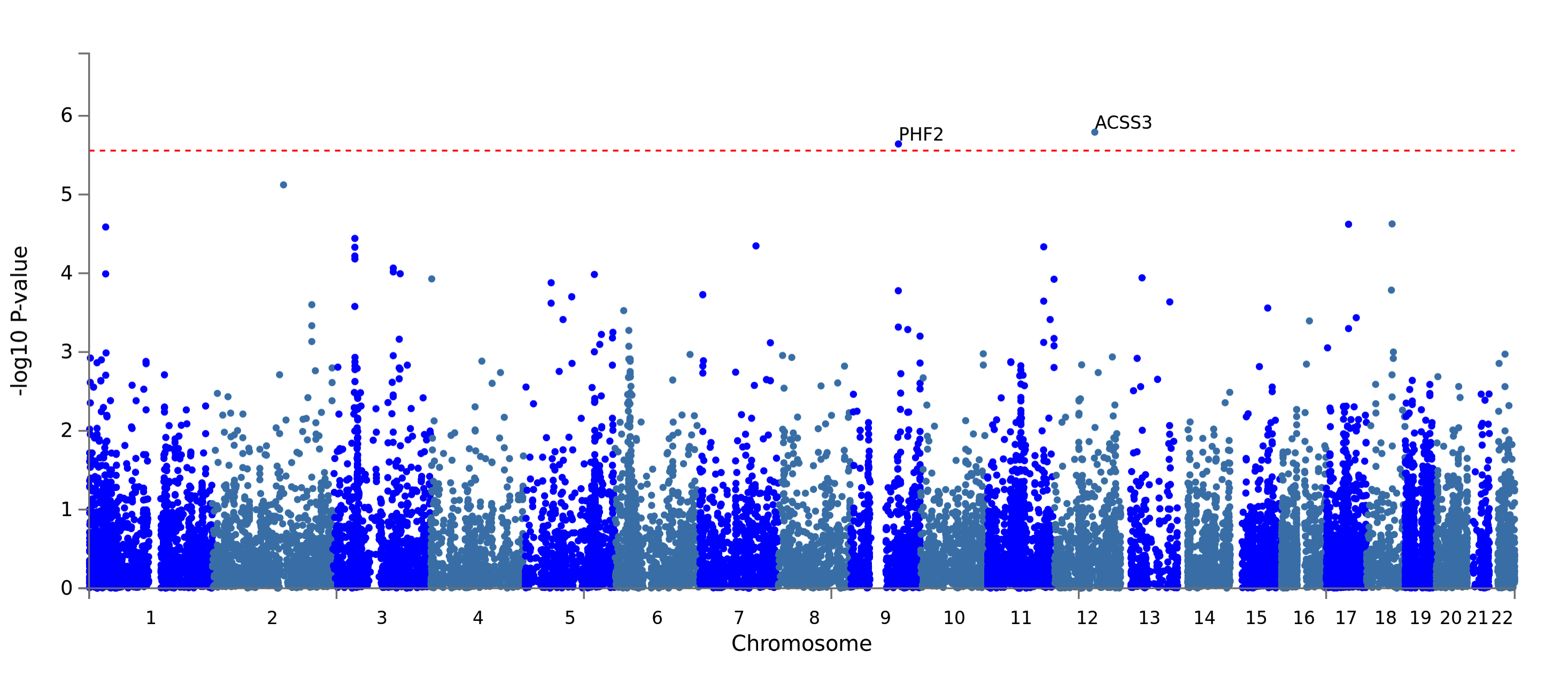

Manhattan plot showing gene-based association of PHQ using summary statistics for the joint effect derived from GWEIS in UK Biobank (N = 99,057). The x-axis is base-paired chromosomal position and the y-axis represents the significance (-log_10_ *p* value) of association between joint effects and PHQ. Bonferroni-corrected significance threshold (*p* = 0.05 / 18,068 = 2.77x10^-6^) is shown by dashed red line.

#### Supplementary Figure 12. Manhattan plot of gene-based test of GHQ GS GWEIS using TSLE as exposure for GxE effect

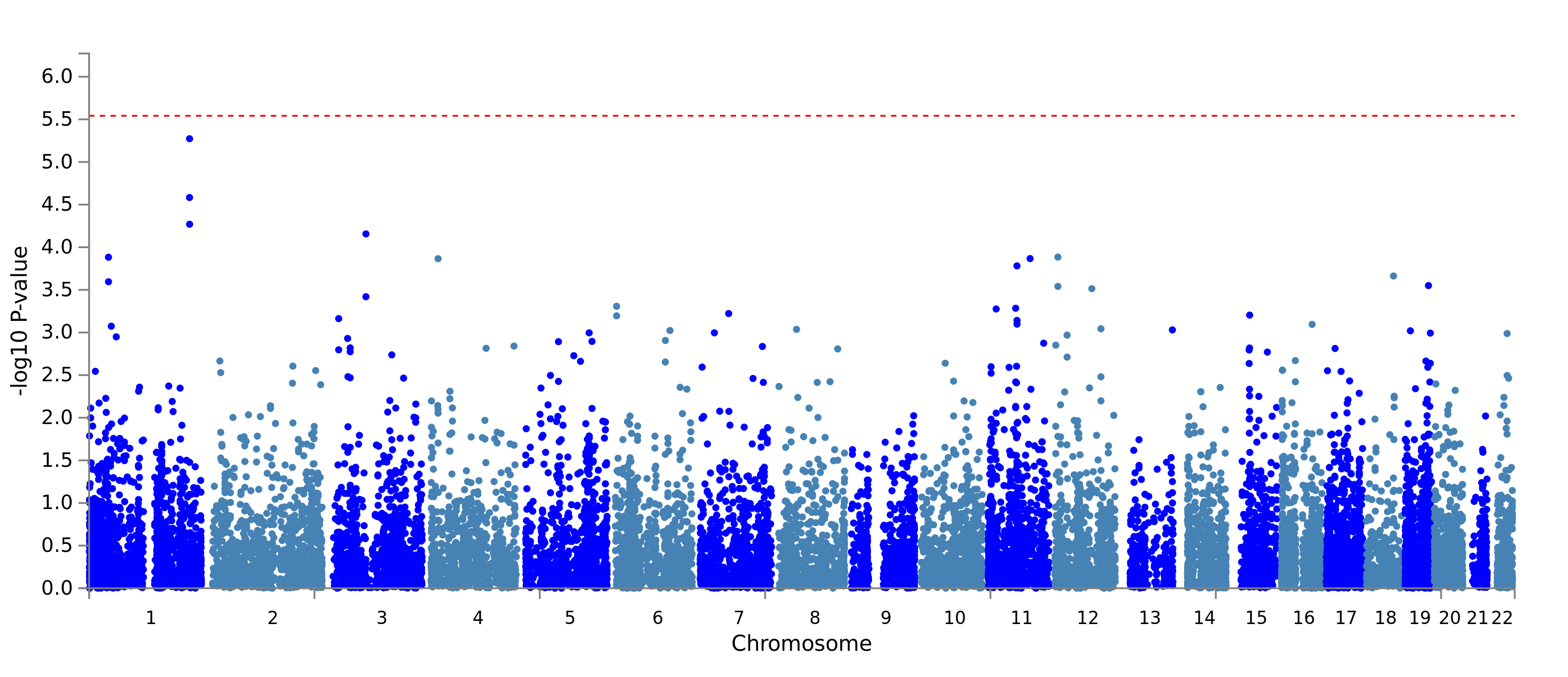

Manhattan plot showing gene-based association of GHQ using summary statistics for the GxE effect derived from GWEIS in Generation Scotland (N = 4,919) using TSLE as exposure. The x-axis is base-paired chromosomal position and the y-axis represents the significance (-log_10_ *p* value) of association between GxE effects and GHQ. Bonferroni-corrected significance threshold (*p* = 0.05 / 18,068 = 2.77x10^-6^) is shown by dashed red line.

#### Supplementary Figure 13. Manhattan plot of gene-based test of GHQ GS GWEIS using TSLE as exposure for joint effect

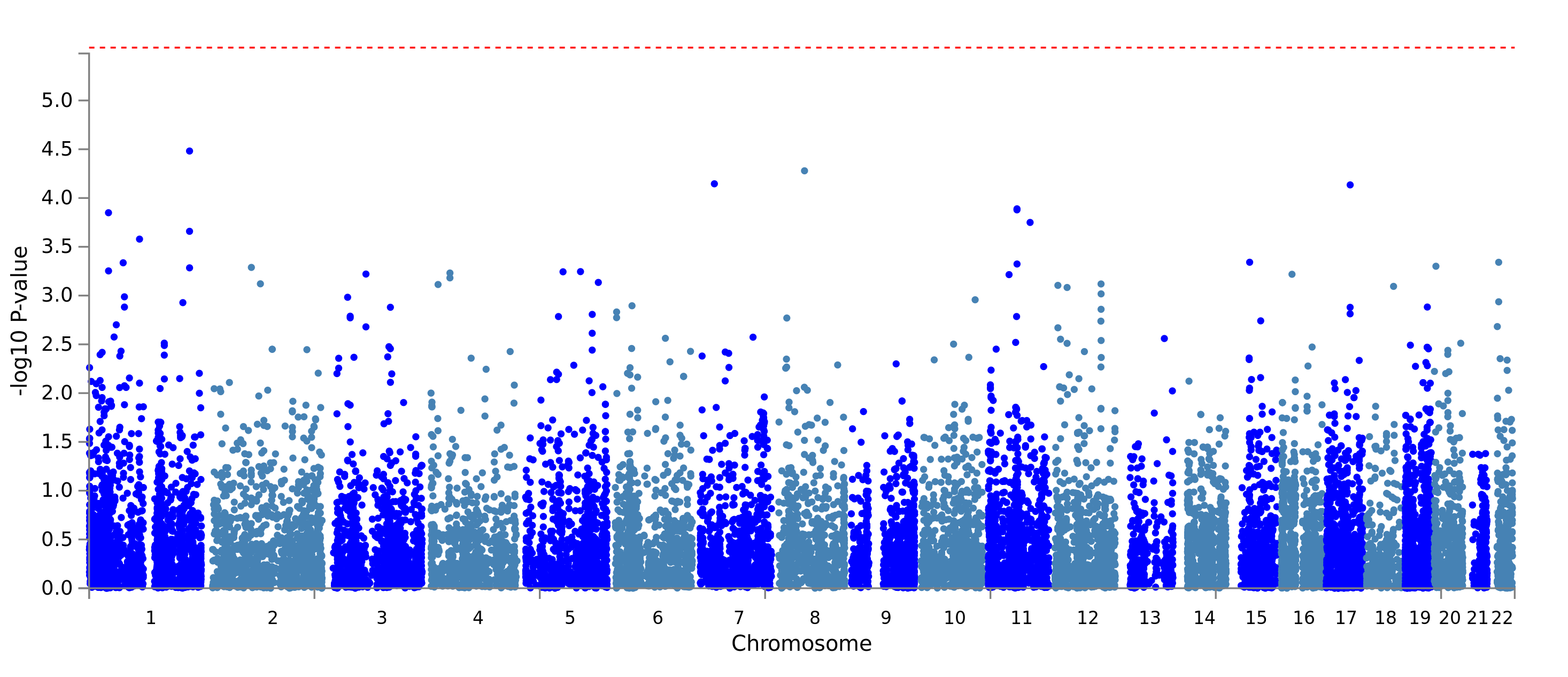

Manhattan plot showing gene-based association of GHQ using summary statistics for the joint effect derived from GWEIS in Generation Scotland (N = 4,919) using TSLE as exposure. The x-axis is base-paired chromosomal position and the y-axis represents the significance (-log_10_ *p* value) of association between joint effects and GHQ. Bonferroni-corrected significance threshold (*p* = 0.05 / 18,068 = 2.77x10^-6^) is shown by dashed red line.

#### Supplementary Figure 14. Manhattan plot of gene-based test of GHQ GS GWEIS using DSLE as exposure for GxE effect

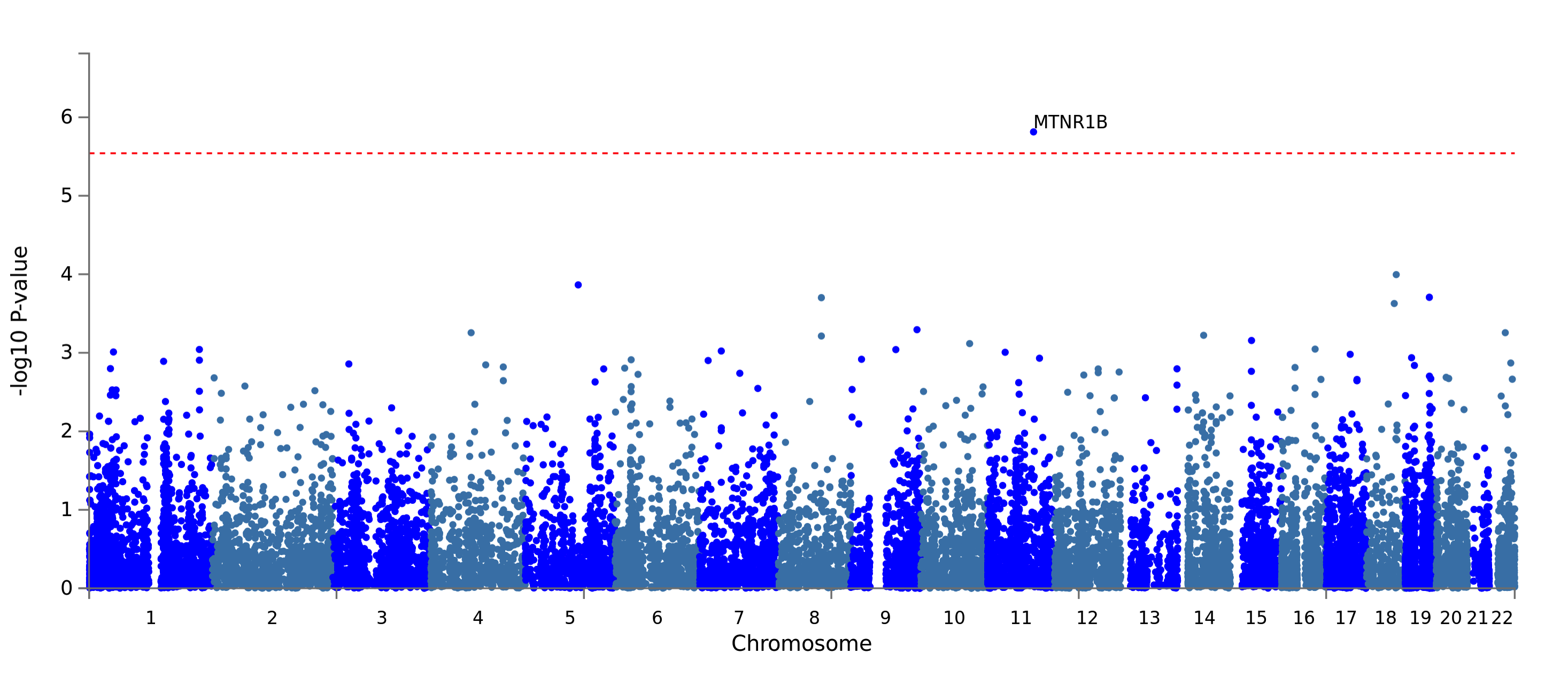

Manhattan plot showing gene-based association of GHQ using summary statistics for the GxE effect derived from GWEIS in Generation Scotland (N = 4,919) using DSLE as exposure. The x-axis is base-paired chromosomal position and the y-axis represents the significance (-log_10_ *p* value) of association between GxE effects and GHQ. Bonferroni-corrected significance threshold (*p* = 0.05 / 18,068 = 2.77x10^-6^) is shown by dashed red line.

#### Supplementary Figure 15. Manhattan plot of gene-based test of GHQ GS GWEIS using DSLE as exposure for joint effect

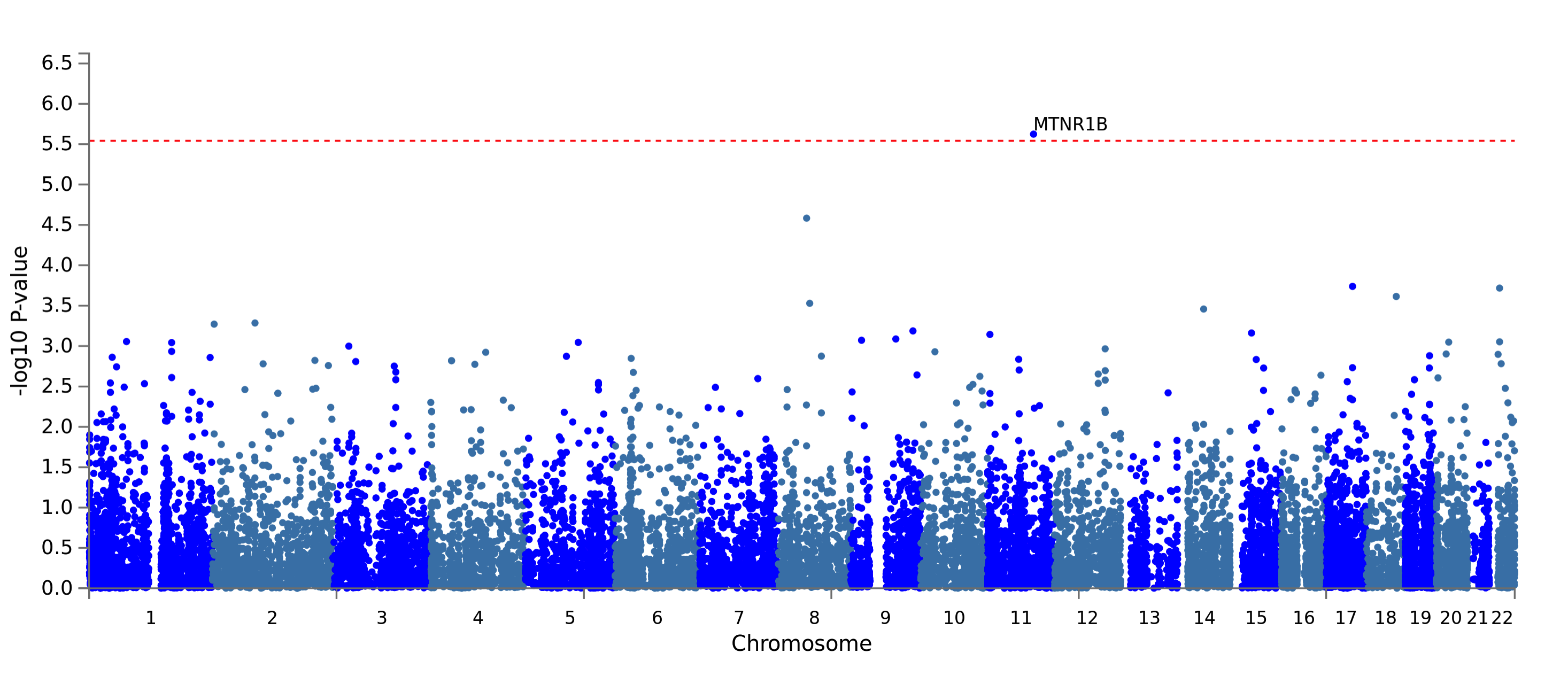

Manhattan plot showing gene-based association of GHQ using summary statistics for the joint effect derived from GWEIS in Generation Scotland using DSLE as exposure. The x-axis is base-paired chromosomal position and the y-axis represents the significance (-log_10_ *p* value) of association between joint effects and GHQ. Bonferroni-corrected significance threshold (*p* = 0.05 / 18,068 = 2.77x10^-6^) is shown by dashed red line.

#### Supplementary Figure 16. Manhattan plot of gene-based test of GHQ GS GWEIS using ISLE as exposure for GxE effect

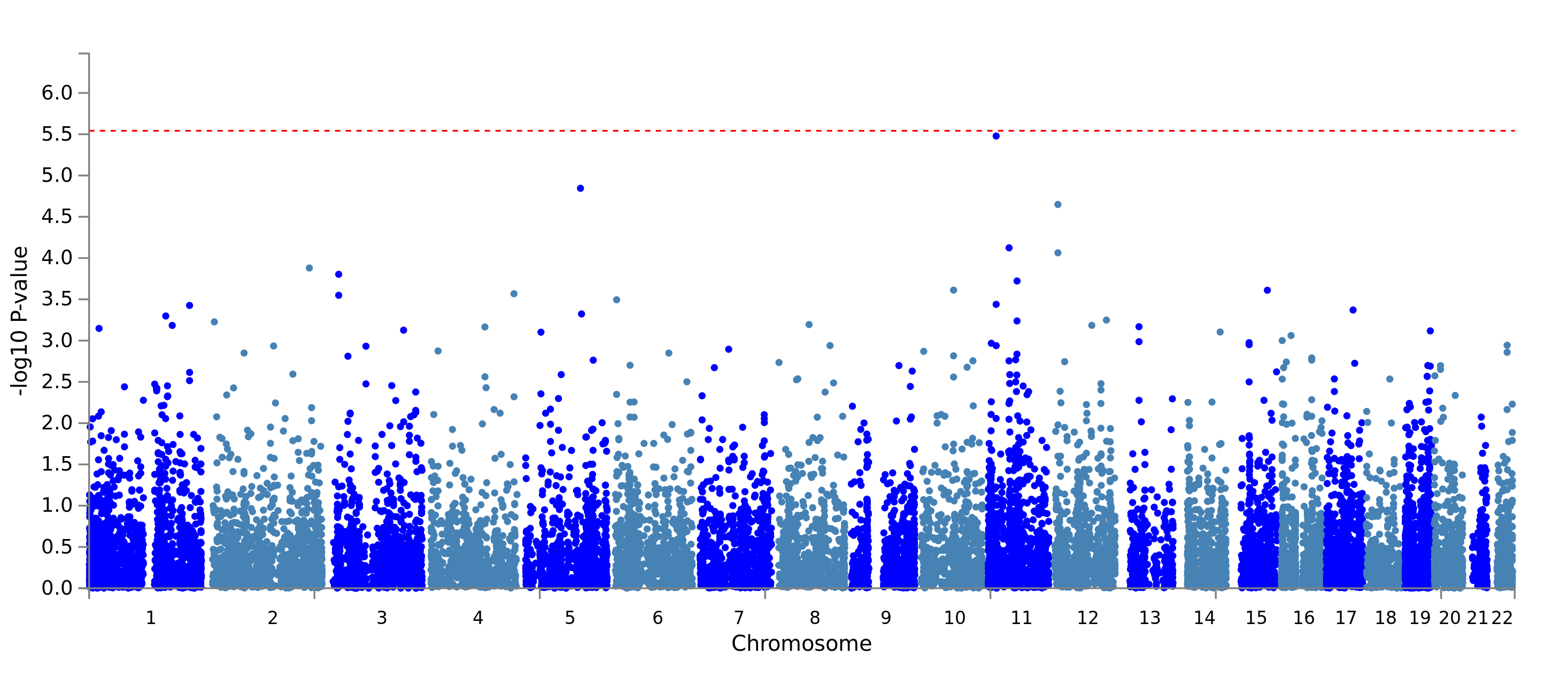

Manhattan plot showing gene-based association of GHQ using summary statistics for the joint effect derived from GWEIS in Generation Scotland (N = 4,919) using ISLE as exposure. The x-axis is base-paired chromosomal position and the y-axis represents the significance (-log_10_ *p* value) of association between joint effects and GHQ. Bonferroni-corrected significance threshold (*p* = 0.05 / 18,068 = 2.77x10^-6^) is shown by dashed red line.

#### Supplementary Figure 17. Manhattan plot of gene-based test of GHQ GS GWEIS using ISLE as exposure for joint effect

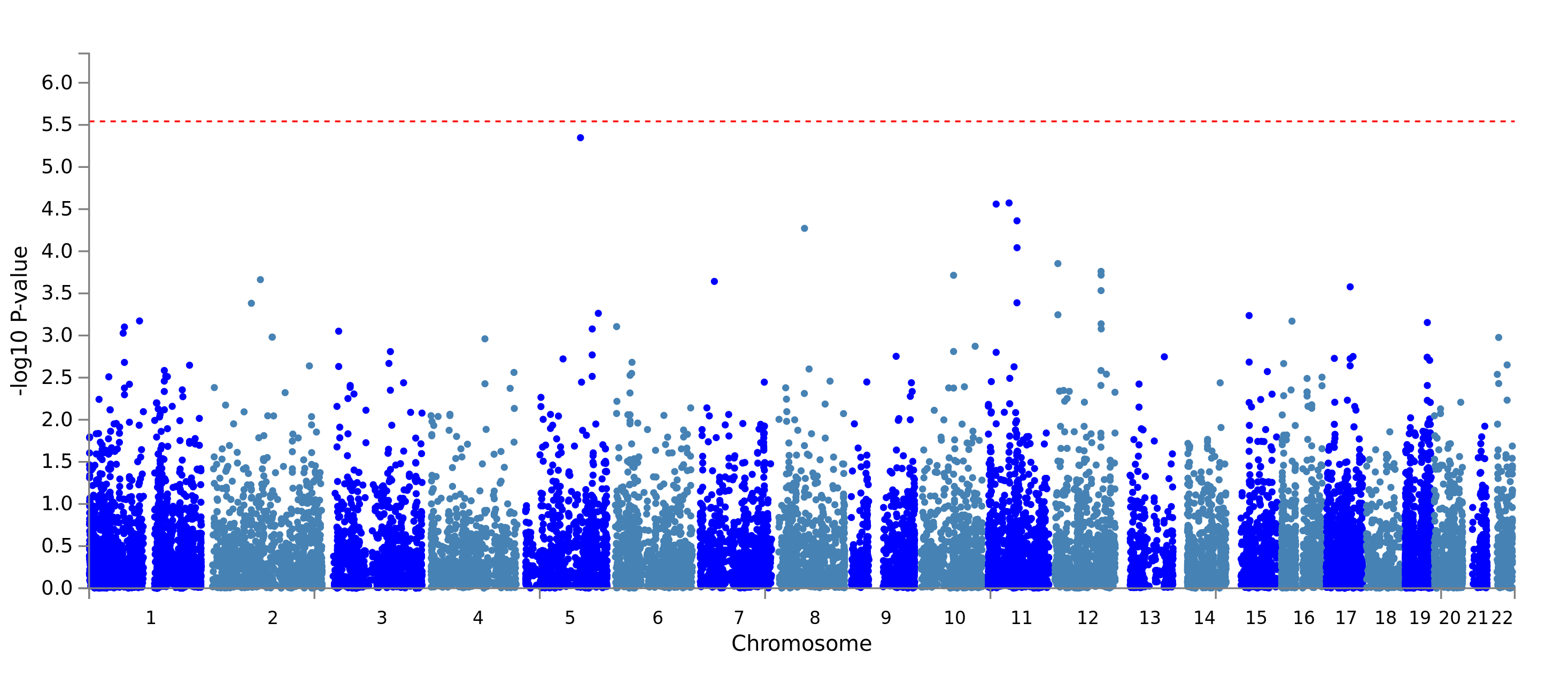

Manhattan plot showing gene-based association of GHQ using summary statistics for the joint effect derived from GWEIS in Generation Scotland (N = 4,919) using ISLE as exposure. The x-axis is base-paired chromosomal position and the y-axis represents the significance (-log_10_ *p* value) of association between joint effects and GHQ. Bonferroni-corrected significance threshold (*p* = 0.05 / 18,068 = 2.77x10^-6^) is shown by dashed red line.
